# Chronic Inflammation Induced Immature Neutrophils Drive Immunopathology During Subsequent Inflammatory Events

**DOI:** 10.1101/2025.04.24.650544

**Authors:** Alakesh Alakesh, Vinod Kumar Dorai, Ranjitha Guttapadu, Shruthi Ksheera Sagar, Meghana Valakatte, Jayashree Vijaya Raghavan, Monisha Mohandas, S R Kalpana, Nagasuma Chandra, Siddharth Jhunjhunwala

## Abstract

Individuals with underlying chronic inflammatory conditions are prone to increased morbidity when posed with an additional inflammatory challenge such as an injury or infection, but why this occurs remains unclear. Herein, we address this question in mouse models by demonstrating that chronic inflammation results in a pronounced expansion of circulating immature neutrophils that exhibit dysregulated effector functions as determined by single cell RNA sequencing and *ex vivo* functional assays. We show that these immature neutrophils are associated with and contribute to increased immunopathology in response to a new inflammatory challenge. Blocking the migration or function of these immature neutrophils through the administration of therapeutic antibodies or small molecules, profoundly lowers the immunopathology caused by the inflammatory challenge. Together, these studies establish a causal link between immature neutrophils and increased immunopathology, while also providing insights into new therapeutic strategies for treating individuals with chronic inflammatory ailments.

## Introduction

Individuals with ailments that are associated with chronic inflammation, like type-2 diabetes mellitus^1^, are prone to increased morbidity and mortality when a second inflammatory event such as an infection^2,3^ or injury^4,5^ occurs. A notable recent example is the case of poor outcomes following SARS-CoV-2 infection in individuals with diabetes and obesity^6^. While the adverse outcomes have been linked generally to aberrant systemic immune responses^6^, such as the increased presence of inflammatory cytokines^7,8^ and expansion of immature and mature myeloid cells during chronic inflammation^9–13^, the specifics are poorly delineated.

Among the many changes to the immune system during chronic inflammation, recent evidence suggests that neutrophil phenotype and function are dramatically altered^14–16^. Specifically, an increase in immature neutrophils (characterized as Ly6G^+^ CD101^-^ in mice^17^ and CD10^-^ in humans^18^) is observed in conditions such as cancer^17^, obesity^19^ or diabetes^20^. Chronic inflammation-driven increase in immature neutrophils has been described to be a result of augmented demand for these cells, resulting in emergency granulopoiesis^16,21^ and altered maturation kinetics^22^. Importantly, higher numbers of immature neutrophils are associated with adverse outcomes in cancer^17^, COVID-19^23,24^ and cardiovascular conditions^25,26^. However, whether and how immature neutrophils contribute to adverse outcomes upon a second insult such as an infection or injury remains unclear.

A significant challenge in addressing this question has been in establishing a pre-clinical model where immature neutrophils are specifically eliminated or increased in large numbers. To address this challenge, previously, we have developed a mouse model involving the implantation of specific biomaterials to induce a dramatic increase in circulating immature neutrophils^22,27,28^. Herein, we show that this model results in changes among neutrophils that are like those observed in leptin-receptor knockout diabetic mice. Then, using the same model, we demonstrate that a specific increase in circulating immature neutrophils drives immunopathology upon a second inflammatory stimulus.

## Results

### Sterile Lung Injury in Obese-Diabetic Mice Shows Greater Immunopathology

Type-2 diabetes has been reported to be a condition that induces chronic inflammation^1^, and hence, we used a mouse model that replicates this condition. Previously, we have reported that leptin receptor knockout mice (LepR KO) that are obese and diabetic show increased numbers and percentages of immature neutrophils (characterized as Ly6G^+^ CD101^-^) in circulation^20^. We confirm these findings (Suppl. Fig. 1A-C) and show that in ex vivo cultures, the circulating immune cells in LepR KO mice produce significantly greater levels of reactive oxygen species (ROS, Suppl. Fig. 1D) and extracellular DNA (a neutrophil extracellular trap (NET)–like action) upon stimulation (Suppl. Fig. 1E) but have lower phagocytic ability (Suppl. Fig. 1F) as compared to their wild-type littermate controls. These data suggest that circulating neutrophils and possibly other immune cells are functionally altered in the LepR KO mice.

In these obese-diabetic mice, when a sterile inflammatory injury is introduced through intra-tracheal administration of lipopolysaccharide (LPS) at a concentration of 0.4mg/kg (amount/body weight of mouse), we demonstrate that there is increased lung injury using weighted histology grading (Suppl. Fig. 2A), resulting from presence of neutrophils in the interstitial spaces and increased hyaline membrane formation (Figure 1A-C). In addition, we observe a significant decrease in the stiffness (Figure 1D and 1E, mean Young’s modulus = 1563 Pascals vs. 427 Pascals for WT and LepR KO, respectively) and hydroxyproline content (Figure 1F) of the lung tissue, possibly due to greater immune cell infiltration and damage caused by such infiltration, in the LepR KO mice as compared to wild-type littermate controls. The baseline stiffness of lung tissue was similar in these mice (Suppl. Fig. 2B). We also observed increased mortality among the LepR KO mice when a higher dose (2mg/kg) of LPS was administered (Suppl. Fig. 2C). Together, these data suggest that in the LepR KO mice the circulating neutrophils have altered function, and when a second inflammatory stimulus (LPS-mediated sterile lung injury) is introduced, it results in adverse outcomes.

**Figure 1:**
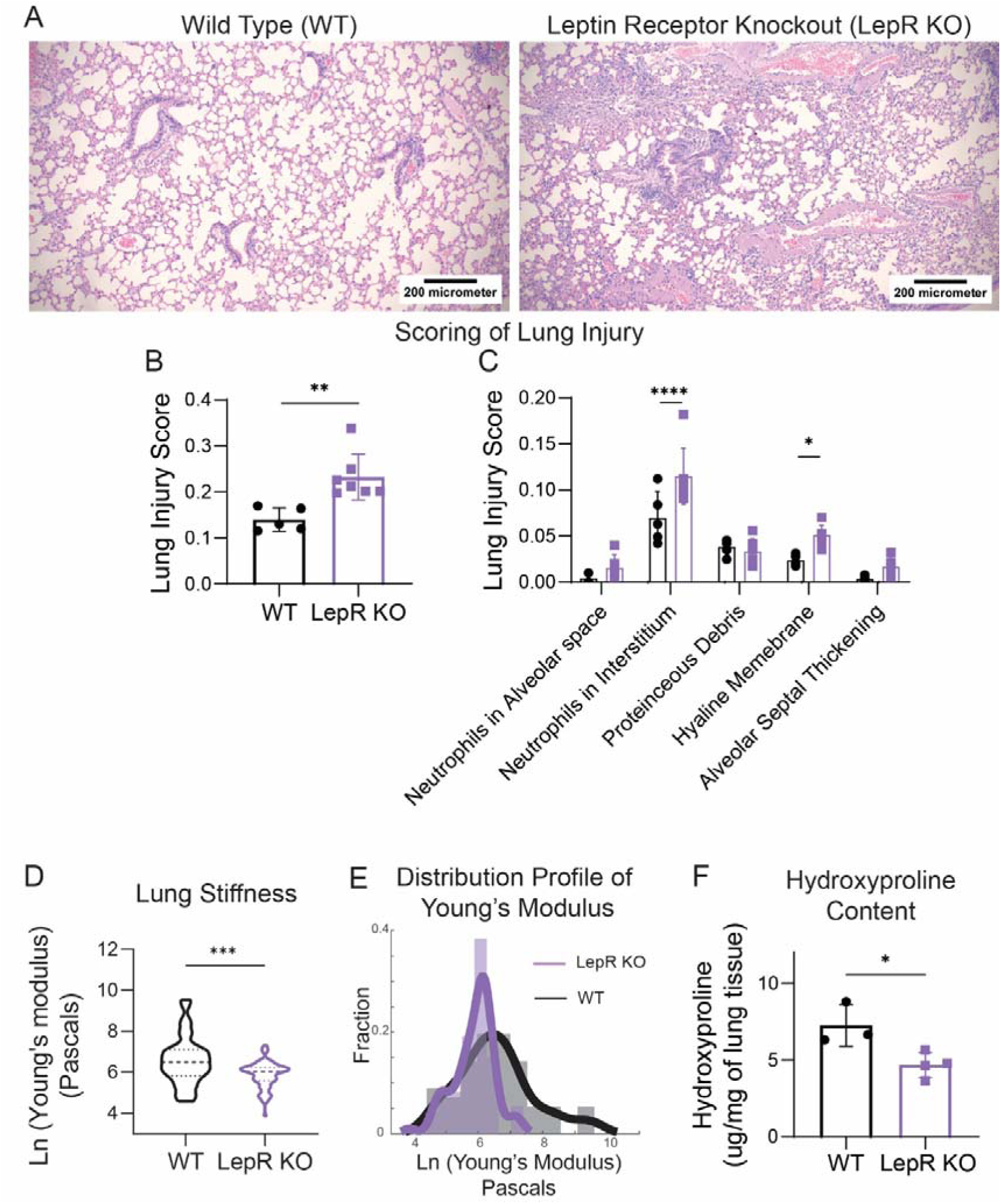
Sterile lung injury in obese-diabetic leptin receptor knockout (LepR KO) mice shows greater immunopathology. **A** – Representative images of hematoxylin and eosin-stained tissue sections at 28 days post lipopolysaccharide (LPS) administration. scale bar = 100µm. **B and C** – Lung damage was quantified from these images and represented as (B) overall injury score and (C) a breakdown of individual weighted scores from each parameter used for overall scoring of the images. For data presented in A-C, n=5 animals for wild type (WT) and 7 animals for LepR KO. **D** – Lung stiffness measured by micro-indentation using atomic force microscopy, and **E** – the distribution profile of Young’s modulus obtained from each measurement (dark lines indicate curves fitted on the histograms using a probability distribution fit). N=56 points for WT and N=73 points for LepR KO micro indentations sampled from n≥3 animals of each strain. F – Hydroxyproline content measured using chloramine T assay; n=3 for WT and n=4 for LepR KO. Student’s t-test or a two-way ANOVA was used to analyze statistical significance among data presented in this figure, with * representing p<0.05, ** p<0.01, *** p<0.001, and **** p<0.0001.

However, the two observations; that there are increased circulating immature neutrophils, and that the injury score is significantly higher upon sterile lung injury in the obese-diabetic LepR KO mice, are correlative. This is especially true in obese-diabetic mice, where multiple cell types are altered. To examine the link between immature neutrophils and increased immunopathology, we needed a model that exclusively has increased or decreased immature neutrophils in circulation, with minimal alterations in other cell types.

### Biomaterial-Implant Model Results in Increased Circulating Immature Neutrophils with Distinct Functional Changes

Previously, we have demonstrated that the implantation of specific sterile biomaterials at a local site in the body induces systemic changes in neutrophil kinetics along with a prominent increase in immature neutrophils in circulation^22^. In this model of chitosan implantation (referred to as chitosan henceforth), we confirm that ∼80% of the circulating neutrophils are immature (Suppl. Fig. 3A and 3B), and demonstrate that in ex vivo cultures, the circulating immune cells have altered function such as increased ROS (Suppl. Fig. 3C) and extracellular DNA (Suppl. Fig. 3D) secretion but lowered phagocytic ability (Suppl. Fig. 3E). These functional changes in circulating neutrophils and other immune cells resemble the changes observed in obese-diabetic LepR KO mice.

To validate the phenotype of these immature neutrophils and confirm that there are no major changes among other cell types (apart from those in monocytes that are likely associated with changes in neutrophils), we performed single cell RNA sequencing (scRNAseq) of circulating cells from mice with chitosan implants and controls (mock surgeries with no implants). Upon clustering and annotation of major immune cell types (Suppl. Fig. 4A), we observed that the neutrophils were proportionally higher in the chitosan-implanted mice (Figure 2A), which correlates with our counts and percentages data in Suppl. Fig. 3B and published previously^22^. Apart from the increased proportion of neutrophils, we also observed that changes in neutrophils and monocytes accounted for greater than 94% of the differentially expressed genes (DEGs) among the circulating cells (Figure 2B). Further, the proportion of neutrophils lacking CD101 (immature neutrophils) gene transcripts was significantly elevated in the biomaterial-implanted mice (Suppl. Fig. 4B) in concordance with our flow cytometry data.

**Figure 2:**
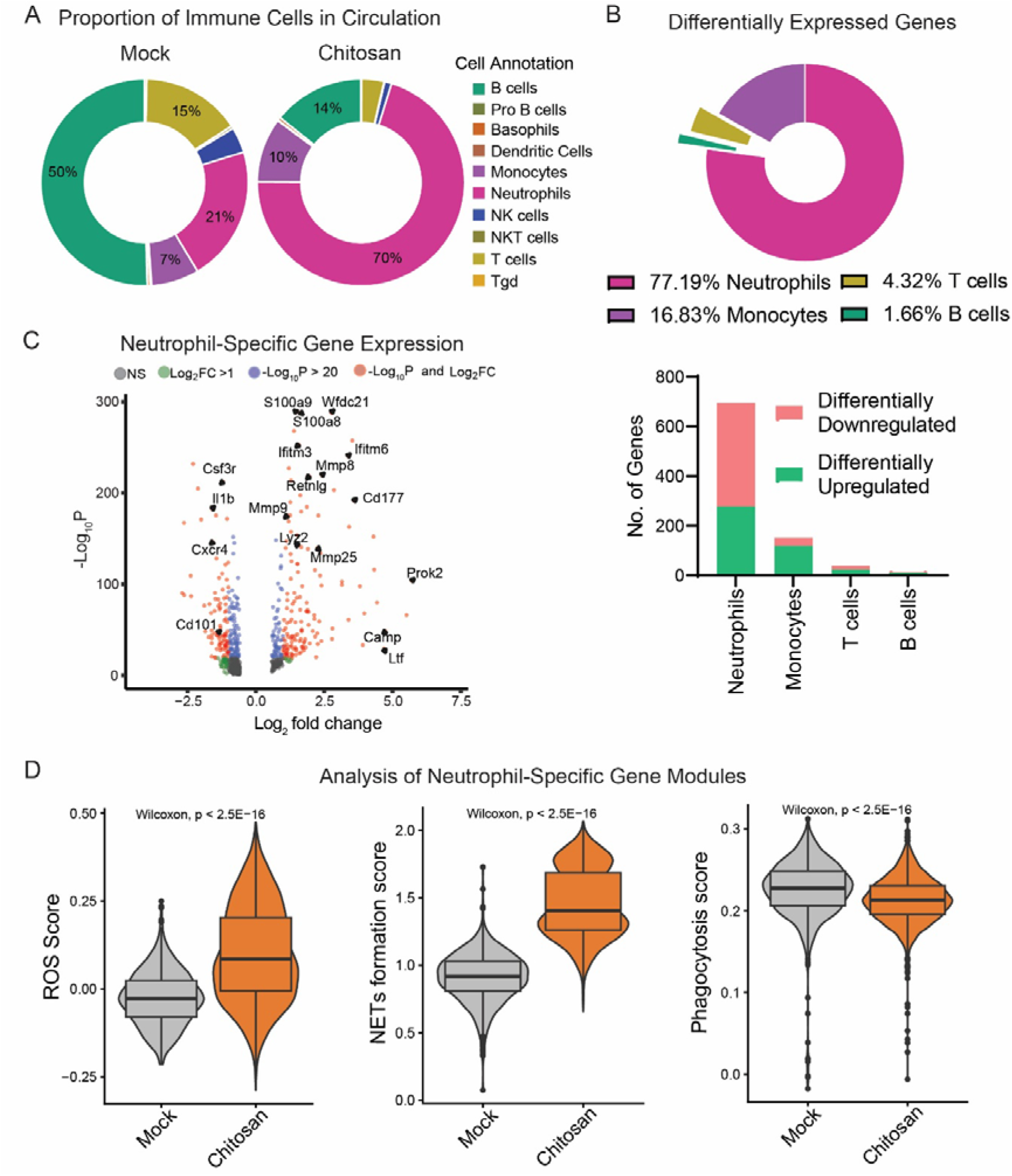
Biomaterial implantation in wild-type (WT) mice results in increased circulating immature neutrophils with functional changes like LepR KO. **A** – Proportions of various immune cells in circulation obtained by analyzing the single cell RNA sequencing (scRNAseq) data of circulating immune cells. **B** – (Top) Measure of the differentially expressed genes (DEGs) among circulating immune cells showing that most of the changes were in neutrophils, and (Bottom) the proportions of DEGs in chitosan-implanted mice (vs. mock). **C** – Enhanced volcano plot of DEGs among neutrophils. **D** – Comparison of neutrophil-specific functional scores (ROS, NETs formation and Phagocytosis score) based on the expression of groups of gene contributing to this function. The significance among functional scores was determined using the Wilcoxon test.

Next, using Ly6G-antibody-derived tags (Ly6G-ADTs), we analysed only the neutrophils in the scRNAseq dataset. The whole neutrophil transcriptome profiling revealed that inflammatory genes (such as MMPs, calprotectin (S100a8 and S100a9), lactotransferrin (Ltf), and Ifitim) were upregulated, while the maturation genes (CD101) were downregulated in the chitosan-implanted mice (Figure 2C). Specifically, the neutrophils showed a significant increase in gene modules that promote ROS production and NET formation, and reduced gene module score for phagocytosis in the biomaterial-implanted mice as compared to mock-treated controls (Figure 2D), which matched our functional data from *ex vivo* cultures (Suppl. Fig. 3C-E). Functionality scores of the granular content, activation status, aging, and positive regulation of apoptosis were found to be different too (Suppl. Fig. 4C), all suggesting an upregulation of an inflammatory neutrophil sub-type.

Then, through fine-resolution UMAP clustering of the Ly6G-ADTs dataset, we identified 5 neutrophil clusters (Figure 3A), among which the i2 cluster was found to be uniquely present in the biomaterial-implanted mice. These clusters were validated by performing a Jaccard similarity analysis using the report by Xie et al.^29^, and the i2 cluster was classified as the one that resembled immature neutrophils or their progenitors (Figure 3B). Upon further investigation, it was observed that the i2 cluster was highly enriched in CD101-negative immature neutrophils, which showed higher ROS, NET formation, NADPH, and activation scores while having lower phagocytosis and maturation scores (Figure 3C).

**Figure 3:**
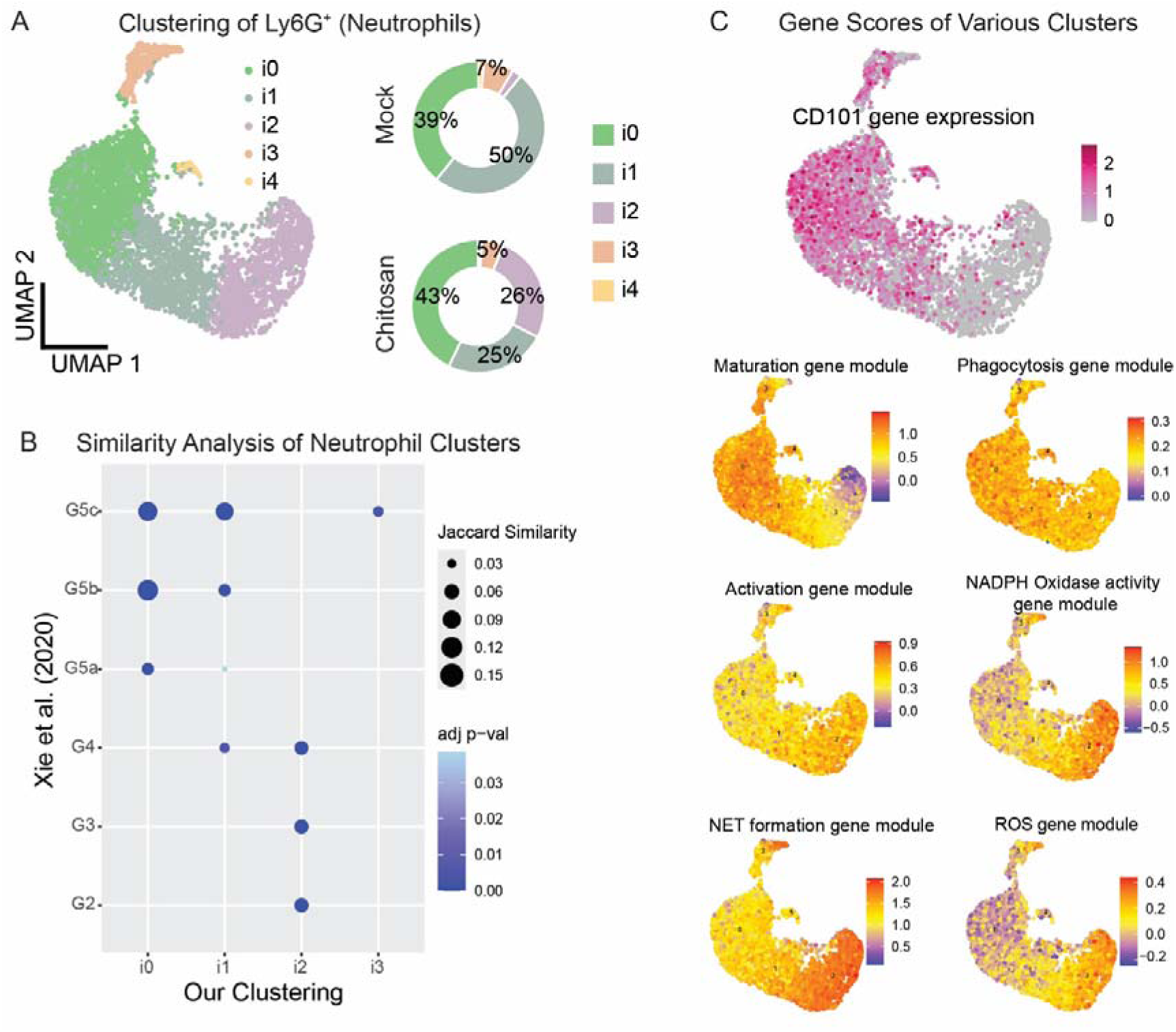
Immature neutrophils expand in chitosan-implanted mice and their transcriptional landscape is associated with functional changes. **A** – Unbiased uniform manifold approximation and projection (UMAP) of neutrophils from mock and chitosan-implanted mice. Each dot represents an individual cell and is colored according to cell cluster. Proportion of each cluster is represented as donut plot. **B** – Pairwise Jaccard similarity analysis of the clusters identified in A with a published scRNAseq dataset. **C** – Feature plot of CD101 gene expression superimposed on neutrophil clusters identified in A suggesting the absence of CD101 expression in i2 cluster, and neutrophil-specific functional scores superimposed across these clusters showing the gene modules that are potentially upregulated or downregulated in these clusters.

Together, these data show that in the chitosan-implanted mice, most of the circulating neutrophils are immature and pro-inflammatory, and other than the neutrophils and monocytes, no other immune cell type showed major changes. Hence, this model could be used to test our hypothesis that an increase in circulating immature neutrophils causes greater immunopathology upon a second inflammatory insult.

### Sterile Lung Injury in Chitosan-Implanted Mice Shows Greater Immunopathology

We induced a sterile lung injury using intratracheal administration of LPS in the chitosan- implanted and control (mock-surgery) mice (Suppl. Fig. 5A). We observed that biomaterial-implanted mice had worse lung injury scores as compared to controls, which was driven by increased neutrophil presence in the alveolar and interstitial spaces at day 14 (Figure 4A, Supp. Fig 5B), and increased interstitial neutrophils with hyaline membrane formation at day 28 (Figure 4B, Supp. Fig. 5C), mirroring lung damage observed in obese-diabetic LepR KO mice (Figure 1C). In addition, we observed a similar shift in the Young’s modulus distribution profile and reduced stiffness of lung tissue in implant bearing mice (Figure 4C-D), with the mean lung stiffness for mice not treated with LPS was measured as 3058 Pascals, while mock-surgery followed by LPS treatment was 1831 and 1837 Pascals (14- and 28-day, respectively), and chitosan-implanted followed by LPS treatment was 660 and 1277 Pascals (14- and 28-day, respectively) .

**Figure 4:**
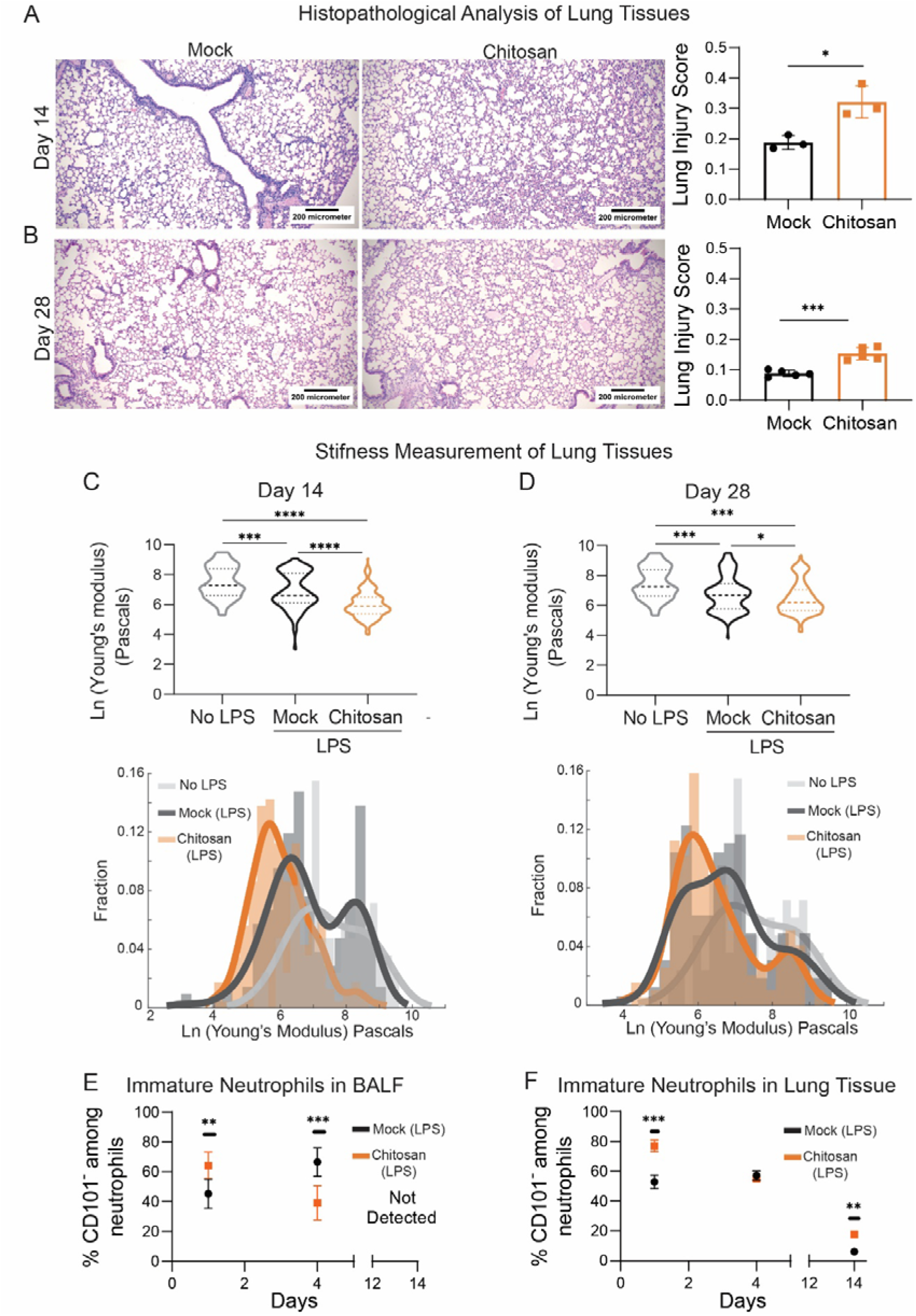
Sterile Lung Injury in Chitosan-Implanted Mice Shows Greater Immunopathology. **A and B** – Representative images of hematoxylin and eosin-stained tissue sections at 14 days post (A) or 28 days post (B) lipopolysaccharide (LPS) administration (scale bar = 100µm) and quantification of lung damage as an injury score (right). n=3 animals/group (A) and n=5 animals/group (B). **C and D** – Lung stiffness measured by micro-indentation using atomic force microscopy (Top), and the distribution profile of Young’s modulus (Bottom) obtained from each measurement (dark lines indicate curves fitted on the histograms using a probability distribution fit) at 14 days post (C) and 28 days post (D) LPS administration. N=210 for mock and N=234 for chitosan micro indentations sampled across n≥3 animals/group (14-days post) and N=163 for mock and N=195 for chitosan micro indentations sampled across n≥3 animals/group (28-days post). No LPS control animals (n=3), N=84 micro-indentations sampled. **E and F** – kinetics of immature neutrophils in bronchoalveolar lavage fluid (BALF, E) and lung tissue (F) assessed using flow cytometry. n ≥ 3 animals/group at each time point. Student’s t-test, one-way ANOVA or two-way ANOVA was used to analyze statistical significance among data presented in this figure, with * representing p<0.05, ** p<0.01, *** p<0.001, and **** p<0.0001.

Associated with these changes in lung injury were alterations in the kinetics of neutrophils and monocytes in the bronchoalveolar lavage fluid (BALF) of these mice measured by flow cytometry. While the total numbers of neutrophils was not dramatically altered in the BALF (Suppl. Fig. 5D), we observed an early presence (1-day post lung injury) of a significantly higher proportion of immature neutrophils with a switch to a significantly lower proportion by 4-day post lung injury, in the bronchoalveolar lavage fluid (BALF) of chitosan-implanted mice as compared to control mice (Figure 4E). Correspondingly, in the lung tissue (Suppl. Fig. 5E) too, we observed an increased early presence of immature neutrophils (Figure 4F). Importantly, an increased and continued presence of immature neutrophils was observed on day 14-post lung injury in the lung tissues using flow cytometry (Figure 4F), which correlated well with our histology data (Suppl. Fig 5B). Accompanying these changes was the early occurrence of a significantly higher proportion of BALF-associated (Suppl. Fig. 5F) and lung tissue-associated (Suppl. Fig. 5G) CD38-expressing monocytes that are thought to be inflammatory^30^.

As the numbers of the cells obtained from BALF and lung tissue are very low (even if we pool cells from multiple mice) to perform ex vivo functional experiments, we resorted to analyzing the potential changes in these cells using scRNAseq of cells obtained from the blood and BALF at day-4 post-injury. This specific time-point was chosen to enable an analysis of neutrophils, monocytes, and macrophages, with the latter two not expected to change immediately (1-day) post-injury. Many differences were noted in the chitosan- implanted mice as compared to the controls.

First, the acute injury caused by LPS administration induces an increase in circulating immature neutrophils in both the mice that have received mock-surgery (prior to LPS) and those with chitosan (Suppl. Fig. 6A-B) as compared to those mice that did not receive LPS (see Suppl. Fig. 3 and 4, mock-surgery groups). Notably, in the chitosan-implanted mice, not only was the proportion of circulating immature neutrophils (assessed using both scRNAseq and flow cytometry) higher than mock-surgery (Suppl. Fig. 6B-C), but these cells also exhibited higher activation, ROS, NADPH and NET scores (Suppl. Fig. 6D). Second, in the BALF, most of the DEGs (>97.5%) were observed among neutrophils, macrophages and monocytes (Figure 5A). Third, in the BALF, in concordance with our flow cytometry data shown earlier (Suppl. Fig. 5E), while the proportion of CD101 lacking immature neutrophils was lower (on day 4) in the chitosan-implanted mice (Figure 5B), they continued to show higher NADPH oxidase activity, NET formation, gelatinase granule and secretory vesicle scores (Figure 5C).

**Figure 5:**
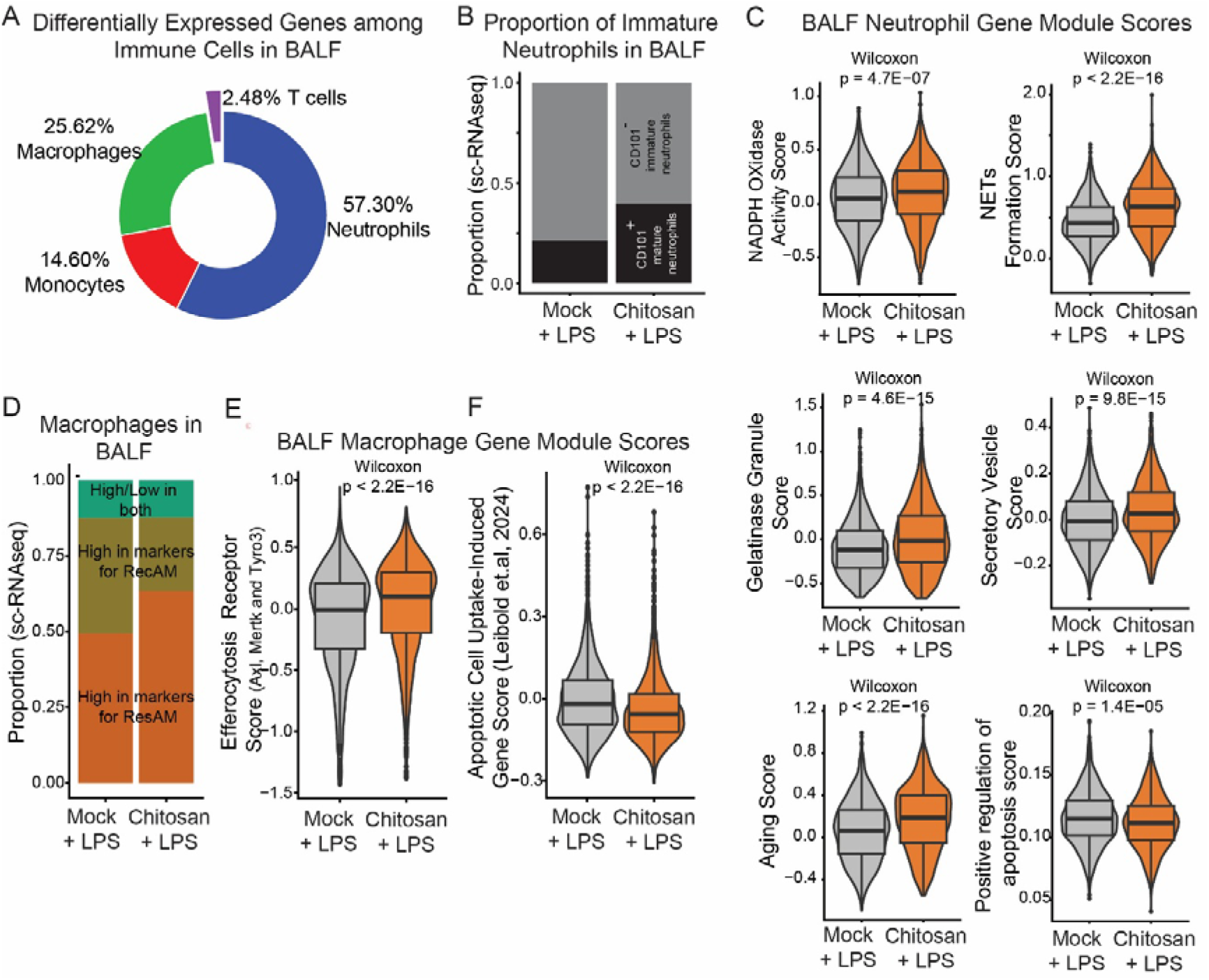
Neutrophils and macrophages in bronchoalveolar lavage fluid 4-days post lipopolysaccharide administration in chitosan-implanted mice show significant alterations. **A** – The proportion of differentially expressed genes (DEGs) expressed as a function of immune cell type showing that most DEGs were among neutrophils, macrophages and monocytes. **B** – Proportion of immature/mature neutrophils in BALF at 4 days post LPS administration based on the scRNAseq data. **C** – Neutrophil-specific gene module scores among BALF neutrophils suggestive of the function of these cells. **D** – Proportion of macrophages of different origin based on their gene signatures (ResAM-Resident Airspace Macrophage, RecAM – Recruited Airspace Macrophage). **E and F** – Gene module scores quantified from BALF macrophages for their ability to express three specific efferocytosis receptors (E) and genes induced following apoptotic neutrophil uptake as published in Leibold et al., 2024 (F). The significance between mock and chitosan groups in the gene module scores was determined using the Wilcoxon test.

Fourth, in the BALF, an increase in the proportion of macrophages was observed (Suppl. Fig. 6E). Among the macrophages, a mix of resident airspace macrophages (ResAM) and recruited airspace macrophages (RecAM) were observed, as is expected following sterile injury with LPS^31,32^. The proportions of ResAM and RecAM followed the expected kinetics (almost equal proportions at day 4 post-LPS injury) in the mice with mock-surgery (Figure 5D). But in the mice with chitosan implants the ResAM were in much higher proportions (Figure 5D), which is supposed to happen at a later time following LPS-induced sterile injury^31^, suggestive of altered kinetics of macrophage responses too. Importantly, while the overall macrophages in the BALF of chitosan-implanted mice expressed higher levels of efferocytic receptors (Axl, Mer, and Tyro3 combined, Figure 5E), the same macrophages showed a significantly lower gene module score for genes that are to be induced following uptake of apoptotic cells (Figure 5F) as described by Liebold et al^33^. The latter data suggests that the macrophages in the BALF of chitosan-implanted mice have not taken up many apoptotic neutrophils, which could be because the neutrophils from these mice have lower signatures of genes that promote apoptosis (Figure 5C).

The above results suggest that immature neutrophils, which are present for longer periods of time at the inflammatory site, that are less apoptotic, show higher NADPH oxidase activity, produce more NETs and induce altered kinetics of monocytes and macrophages in the lung, play a significant role in the development of immunopathology. To confirm that immature neutrophils drive these changes, we performed the following experiments.

### Reducing Immature Neutrophil Number and Activity

First, we used an alternate model to confirm the above findings, which involves acute increases in circulating immature neutrophils. Granulocyte colony stimulating factor (G-CSF) injection in mice has shown to elicit a transient increase in ‘immature-like’ neutrophils in circulation^34–36^. In recombinant murine G-CSF (rmG-CSF) injected mice, we induced sterile lung injury and observed that the immunopathology was higher (Suppl. Fig. 7A-B), and the lung tissue stiffness was reduced (Suppl. Fig. 7C-D) as compared to saline-injected control mice. Data from this model and other published literature^35,36^ suggests that G-CSF and possibly IL-1β are key mediators in increasing neutrophil numbers.

Additionally, CellChat^37^ analysis of our sc-RNAseq data from the BALF showed that while the overall neutrophil communication with other cell types appears to be similar between the mock-treated and chitosan-implanted mice (Suppl. Fig. 8A), when focusing specifically on the CSF and IL-1 signaling networks^36,38^ we observed that neutrophils from the chitosan- implanted mice communicate with many more cell types (Figure 6A). Hence, to reduce the number of circulating immature neutrophils and interfere with the neutrophil communication with other cell types in the chitosan-implant model, we injected anti-G-CSF or anti-IL1β antibodies that would temporarily block the function of these proteins (Suppl. Fig. 8B). Administration of these blocking antibodies significantly lowered lung injury (Figure 6B-C, Suppl. Fig. 8C) and lessened the reduction in lung stiffness (Figure 6D-E) caused due to LPS-mediated sterile injury in chitosan-implanted mice as compared to control isotype antibody injection.

**Figure 6:**
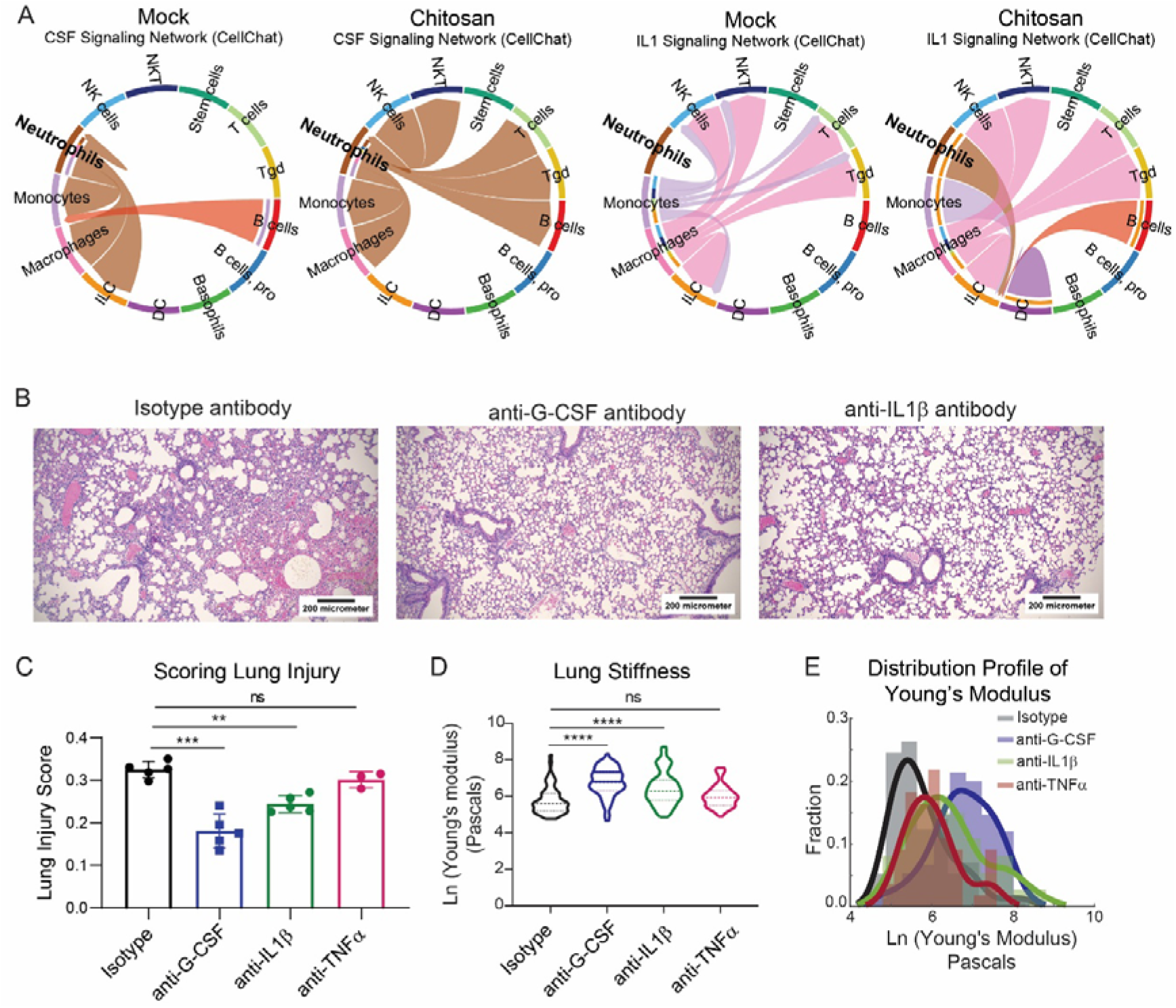
anti-GCSF and anti-IL1β treatment reduces lung damage in chitosan-implanted mice. **A** – Chord plot of CSF and IL1 signaling network analysis using Cellchat performed on immune cells obtained from BALF (4 days post LPS administration). **B** – Representative images of hematoxylin and eosin-stained tissue sections following treatment with anti-GCSF and anti-IL1β, scale bar = 100µm. **C** – quantification of lung damage and its representation as an injury score,) chitosan-implanted mice treated with isotype control antibody (n=5), anti-GCSF (n=5), anti-IL1β (n=5) and anti-TNFα (n=3). **D and E** – Lung stiffness measured by micro-indentation using atomic force microscopy (D), and the distribution profile of Young’s modulus (E) obtained from each measurement (dark lines indicate curves fitted on the histograms using a probability distribution fit). Mice treated with Isotype control (N=57), anti-GCSF(N=75), anti-IL1β(N=74) and anti-TNFα (N=56). For statistical comparisons in panel C and D, a one-way ANOVA followed by Tukey’s post-hoc test was used, and ** represents p<0.01, *** p<0.001, and **** p<0.0001.

In addition, we also tested the effect of anti-TNFα antibody, as TNF-α has a central role in inflammatory responses in the lung following LPS-mediated sterile injury^39,40^. However, administration of anti-TNFα antibody did not alter the lung injury score (Suppl. Fig. 8D and Figure 6C) or change the lung stiffness as compared to the isotype control (Figure 6D-E). Correspondingly, in the CellChat analysis too we did not observe any TNFα related signaling in the chitosan-implanted mice as compared to the mock controls (Suppl. Fig. 8E).

Finally, we tested the effect of sivelestat, which is a neutrophil elastase inhibitor that reduces NET-activity of neutrophils and other inflammatory mediators^41,42^ (Figure 7A). Silvestat administration (in chitosan-implanted mice that subsequently received the LPS-mediated sterile lung injury) showed dramatic effects in preventing immunopathology, with the lung injury score (Figure 7B-C) being significantly lower and lung stiffness (Figure 7D-E) significantly higher than the chitosan-implanted mice (mean Young’s modulus was 735 Pascals for chitosan and 2551 Pascals for sivelestat-treated mice), and the numbers for both these parameters was similar to those of the mock-surgery control mice shown in figure 4. We also observed the proportion of immature neutrophils to be reduced in lung tissue of sivelestat-treated mice (Figure 7F). Together, these data are suggestive of comprehensive prevention of the pathology when one of the activities (NET formation) of immature neutrophils is blocked.

**Figure 7:**
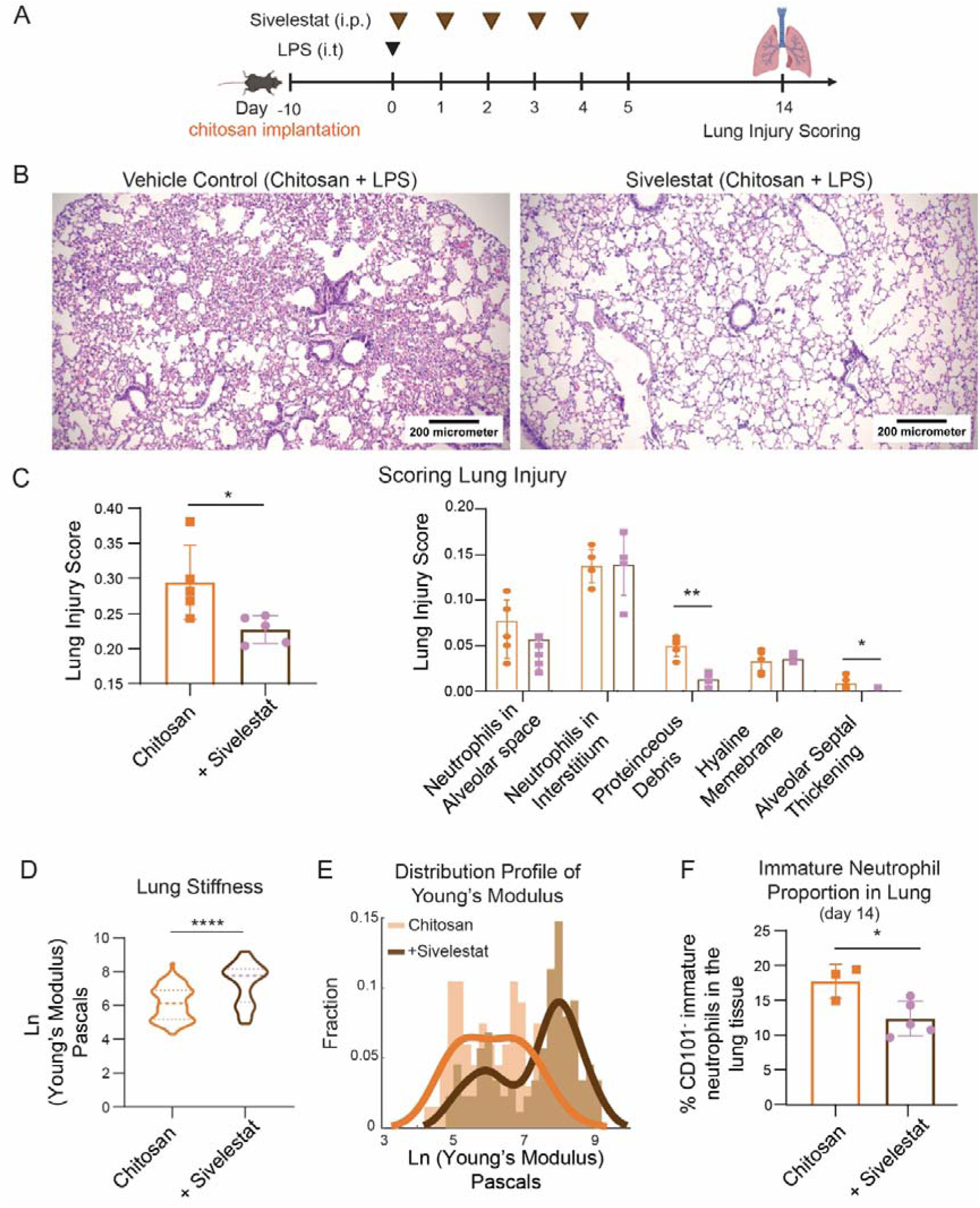
Sivelestat treatment reduces lung damage in chitosan-implanted mice. **A** – Study design for treatment with Sivelestat (50mg/kg, given intraperitoneally (i. p.)) in chitosan-Implanted mice. **B** – Representative images of hematoxylin and eosin-stained tissue sections following treatment with Sivelestat, scale bar = 100µm. **C** – Lung damage was quantified from these images and represented as (left) overall injury score and (right) a breakdown of individual weighted scores from each parameter used for overall scoring of the images. For data presented in B-C n=5 animals/group. **D and E** – Lung stiffness measured by micro-indentation using atomic force microscopy (D), and the distribution profile of Young’s modulus (E) obtained from each measurement (dark lines indicate curves fitted on the histograms using a probability distribution fit). chitosan-implanted mice (N=67) and chitosan implanted mice treated with Sivelestat (N=88). (N - represents the number of micro indentations sampled across n≥3 animals/group). **F** – Percentage of Immature neutrophils (CD101-) quantified using flow cytometry of cells from the lung tissue (14 days post LPS administration).

## Discussion

The COVID-19 pandemic highlighted a well-known fact that individuals with conditions such as diabetes (co-morbidities), which are associated with chronic inflammation, have a significantly higher risk of exaggerated host response following another inflammatory stimulus^43–47^. Indeed, one of the treatments suggested to prevent/treat acute respiratory distress syndrome following SARS-CoV2 infection was the use of pan-immunosuppressive agents that would reduce the risk of lung immunopathology caused due to excessive and uncontrolled inflammation^48,49^. However, the use of such immunosuppressive agents comes with its own risk of worsening the viral infection, increasing the incidence of other bacterial infections, or causing other issues^50,51^. Instead, if the specific features that promote immunopathology could be identified, then blocking just those cells or pathways could emerge as a method to treat individuals with chronic inflammatory conditions. One such feature is the increased presence of circulating immature neutrophils in individuals and mice with chronic inflammation in conditions such as diabetes.

A causative relationship between a specific entity and an outcome may be determined by the addition of it to a system that lacks it or the removal of this entity from the system and then measuring changes in the outcome. In this specific case, our premise was that the entity is immature neutrophils and the outcome they cause is excessive immunopathology. The addition of immature neutrophils into a wild-type mouse that has relatively low numbers of these cells poses many challenges. Traditional methods of addition of cells involve adoptive transfer. But in the case of immature neutrophils, this has not been possible due to difficulties in handling them ex vivo without activating the cells or inducing apoptosis (our experience^28^ and data not shown), the large numbers of cells required to increase their numbers beyond the existing mature neutrophils in a system that are difficult to obtain, and very short life-span post adoptive transfer of neutrophils requiring multiple doses of these cells that is not feasible in such an experimental setup^28,52^. Hence, we chose the method of utilizing biomaterial-implants^27^ that alter the immature neutrophil numbers with minimal changes in other cells (Figures 2 and 3). Using this method, we were able to ensure that most circulating neutrophils were immature, and this condition persisted for at least a few weeks, enabling us to assess the effect of these cells upon a second inflammatory injury. And we demonstrated that increasing immature neutrophils in circulation results in greater immunopathology following a subsequent inflammatory injury.

On the other hand, methods to deplete immature neutrophils or all neutrophils have only been partially successful. Antibody-based depletion (anti-Ly6G treatments in mice) is known to remove mature Ly6G expressing neutrophils from circulation. However, this effect is transient (lasting for a maximum of a few days), and in our opinion, such antibody-based depletion methods remove cells that express that specific receptor but not all neutrophils or their progenitors. Given that the mammalian system is designed to maintain the homeostasis of neutrophils in circulation^53,54^, upon reduction of specific mature neutrophils in circulation, other cells such as progenitors and immature neutrophils lacking that specific receptor take their place^22,28^. Hence, instead of attempting to deplete neutrophils entirely, we utilized two different strategies. One was the use of anti-G-CSF (and anti-IL1β), which reduces the number of neutrophils and has been suggested to reduce neutrophil extravasation^36,38^, which resulted in reduced lung injury as measured 14-days post-LPS administration. The other was to suppress a specific function of immature neutrophils, which is enhanced production of NETs, through the administration of sivelestat^55,56^. Sivelestat dramatically reduced lung immunopathology and associated changes in lung stiffness. Together, these data demonstrated that reducing immature neutrophil numbers or their function in settings with multiple inflammation, results in less immunopathology.

## Limitations of the Study

One of the key limitations of the current study is the inability to remove neutrophils, specifically immature neutrophils, completely from an in vivo system. Neutrophil deficient mouse strains such as the G-CSFR^-/-^ knockout or LysM-Cre Mcl-1^fl/fl^ could be used but were not available to us. While such studies would likely have helped us answer the scientific question of what happens when these cells are entirely removed from the system, our current experimental strategies are possibly closer to real-world scenarios where an increase in immature neutrophils is observed under inflammatory conditions and reducing them using therapeutics could be a viable strategy in the clinic. Another limitation is the lack of ability to measure functional changes in lung tissue. Our assessment of lung injury and damage relies on histopathological analysis and lung stiffness measurements on tissues obtained from euthanized animals. Measurement of lung function, such as pulmonary function tests, in live animals would have provided a better assessment of the real extent of damage. This would be the focus of future studies, especially in the context of therapeutic treatment strategies that can reduce damage associated with inflammatory injury and promote better quality of life in individuals with pre-existing inflammatory conditions.

## Conclusion

In this study, we demonstrate that implantation of biomaterials in mice may be used as a model to replicate the characteristic of increased immature neutrophils in circulation, which occur in conditions associated with chronic inflammation like type-2 diabetes. Using a combination of flow cytometry based cellular analysis, single cell RNA sequencing of circulating immune cells, and ex vivo functional assays, we show that the immature neutrophils are alternately activated, produce higher amounts of reactive oxygen species and neutrophil extracellular trap components, and exhibit reduced rates of apoptosis. Next, through a sterile lung injury model following the implantation of biomaterials, we determine that increases in circulating immature neutrophils caused by the first inflammation are associated with greater immunopathology due to the second inflammatory event. By specifically blocking G-CSF or IL-1β signaling and by blocking neutrophil elastase activity using the small molecule sivelestat, we establish that immature neutrophils specifically drive immunopathology upon a second inflammatory injury. These findings highlight a potential mechanism through which immature neutrophils may drive inflammation and tissue damage, and provide clues for the clinical prevention of such tissue damage.

## Materials and Methods

### Statement on ethics for animal experiments

All animal studies were conducted following the rules and guidelines of the Control and Supervision Rules, 1998, of the Ministry of Environment and Forest Act (Government of India), and the Institutional Animal Ethics Committee, IISc. Experiments were approved by the Committee for Purpose and Control and Supervision of Experiments on Animals at IISc (permit numbers CAF/ethics/718/2019 and CAF/ethics/890/2021).

### Mouse strains

Female C57BL6 mice that were found to be specific pathogen free were purchased from Hylasco Biotechnology (India) Pvt. Ltd, Hyderabad, India (a Charles River Laboratory licensed supplier). The obese-diabetic leptin receptor knockout (LepR KO) mouse strain - B6.BKS(D)-Leprdb/J - was purchased from The Jackson Laboratory (Bar Harbor, Maine, USA.) and were bred and housed at a clean-room facility, in the Central Animal Facility (CAF), Indian Institute of Science (IISc). All the animals were maintained in individually ventilated caging systems (IVCs) on irradiated chow diet (Altromin, Germany), and autoclaved water obtained through reverse osmosis.

### Chitosan Microspheres and Chronic Inflammation mouse model

Chitosan microspheres (337± 47 micrometers) were prepared using the Spraybase® Electrospray system inside a sterile biosafety cabinet and chemically sterilized as described^22^. Microspheres were surgically implanted into the peritoneal cavity of mice to induce chronic inflammation. In brief, microspheres (450 µL) were suspended in 450 µL sterile saline for implantation. The entire volume (900 µL) was then implanted into the peritoneal space^22,27^ following a laparotomy procedure. In controls (also called mock-surgery), the animals went through the surgical procedure and were injected with 900 µL sterile saline.

### Acute Lung injury (ALI) model

Mice were anesthetized with ketamine (120 mg/kg) and xylazine (40 mg/kg) and placed on a board at an angle of 50 degrees in a supine position. The tongue was carefully pulled using sterile forceps, and the chest was illuminated using a cold source light to visualize the tracheal opening. IV cannula (22G) was inserted into the trachea. LPS (0.4 mg/kg, Escherichia coli O127:B8; Sigma-Aldrich, Merck) was administered through the cannula in 25 μL of sterile saline using a 200 μL pipette tip. The mice were gently massaged and allowed to recover on a heating pad.

### Cell preparation from mouse tissues

Mice were first anesthetized using high dose ketamine-xylazine solution. After anesthesia, about 500-1000 μL blood was collected through the retro-orbital vein. Following euthanasia, bronchoalveolar lavage fluid (BALF) was collected using the procedure described previously^57^ with minor modifications. A 20-G cannula was inserted into the trachea and fixed with a 93 ligature. Lungs were then lavaged with 1 mL sterile Ice-cold 1X phosphate buffered saline (PBS) two times (total volume 2 mL) to collect BALF. Fluid recovery was >85% each time. Blood and BALF samples were then centrifuged at 400 RCF for 5 mins at 4°C. The supernatant was stored at -80°C and the cell pellet was subjected to RBC lysis in ACK Lysis buffer and resuspended in 1mL PBS for counting using a hemacytometer. The cells were then used for functional and flow cytometry analysis.

Immune cells from lung tissue were isolated as described^58^. The right superior lobe was minced in 1ml of digest solution (RPMI media with 10% FBS, collagenase and DNase) and transferred into 6-well plate with 2mL of additional digest solution. After 60 min. incubation at 37°C, the tissue was dissociated with a 1 mL syringe and filtered through 50-micrometer cell strainer. After centrifugation at 400g for 5min at 4°C, the cell pellet was subjected to RBC lysis in ACK Lysis buffer and resuspended in 1ml PBS for counting using a hemacytometer. The cells were then used for flow cytometry analysis.

### Flow cytometry analysis

Blood, BALF and lung samples were stained for viability using BD Horizon Fixable Viability dye 510 (BD Biosciences, USA). Thereafter blood cells were stained with the following antibodies for 30 min. at 4°C: anti- Ly6G (1A8) and CD45 (30-F11) were purchased from BD Biosciences and CD101 (Moushi101) purchased from ThermoFisher. Clone names are in parentheses.

Lung and BALF cells were stained with the following antibodies for 30 min at 4°C: CD11b (M1/70), Ly6G (1A8), CD45 (30-F11), Ly6C (Al-21), F4/80 (T45-2342), all purchased from BD Biosciences and CD101 (Moushi101) as well as CD38 (90) purchased from ThermoFisher.

All flow cytometry data were collected using BD FACSCelesta (Becton Dickinson, USA) or BD FACSLyric (Becton Dickinson) and analyzed using FlowJo (Tree-Star, USA) or FCS Express (DeNovo Software, USA). For all protocols, appropriate single color (using compensation beads, BD Biosciences) and fluorescence-minus-one (FMO) controls (using cells) were used to compensate the data and gate positive populations, respectively.

### Ex-vivo functional assays

For the following functional assays, cells isolated from blood were split into two groups: stimulated and unstimulated. For stimulation, cells were first activated using 5 µg/mL Cytochalasin B (Sigma-Aldrich) for 7 min. and 5µM fMLP (Sigma-Aldrich) for 30 mins at 37°C. Unstimulated cells were followed through the stimulation process, replacing Cytochalasin B and fMLP with PBS.

#### Extracellular reactive oxygen species (ROS)

Extracellular ROS was quantified using cytochrome C assay. Briefly, 2 x 10^5^ cells/well were added to 4 different groups: unstimulated, stimulated (with PMA), unstimulated with SOD (superoxide dismutase - quencher for extracellular ROS as negative control) and stimulated with SOD. Cells were added into a 24-well plate containing a reaction mixture (HBSS, superoxide dismutase for SOD wells and PBS for other wells), PMA (for stimulated well and PBS for other wells), and cytochrome C (for all 4 wells). Absorbance (at 540, 550, and 560nm) was read every 30 mins, and O^2-^ concentration was mathematically calculated using Beer-Lambert’s law as described previously^59^.

#### Intracellular ROS

For intracellular analysis, 2 x 10^5^ cells were plated in 24-well plate in RPMI 1640 media (containing 10% fetal bovine serum, 1% antibiotic-antimycotic, and 25mM HEPES solution) and incubated in a 37°C incubator for 30 min. Cells were split into unstimulated and stimulated groups. Both the groups received 5µM of DHR123 and incubated in 37°C incubator for 30 min. At the end of incubation, cells were scraped from the wells, and the suspension spun down at 400 RCF for 4 min; the cell pellet was washed and suspended in a staining buffer. Cells were then stained with a monoclonal antibody against CD45 (clone 30-F11) and Ly6G (clone 1A8) for 30 mins at 4°C. Further, cells were re-suspended in PBS containing propidium iodide (PI) at a concentration of 2 µg/mL. Median Fluorescence intensity was quantified on Neutrophils (CD45^+^Ly6G^+^) using flow cytometry.

#### Extracellular DNA (a measure of NETosis)

Cells, 1 x 10^5^, were plated in 24-well plate in Imaging solution (NaCl, Kcl, CaCl2, MgCl2, HEPES and D-Glucose) and placed in a 37°C incubator with 5% CO2 for 15 min. After 15 min., cells were split into two groups: unstimulated and stimulated with 50nM of PMA (PMA was a better activator for extracellular DNA release in our experiments). Cell supernatant post-stimulation or supernatant from unstimulated cells was collected in 96-well black plates and assayed with 5uM SYTOX Green (an extracellular DNA binding cell impermeable dye). Fluorescence was quantified at 480/530nm, quantitating DNA in cell supernatant.

#### Phagocytosis

Cells, 2 x 10^5^, were plated in a 24-well plate in RPMI 1640 media (containing 10% fetal bovine serum, 1% antibiotic-antimycotic, and 25mM HEPES solution) and incubated in a 37°C incubator for 30 min. Following initial incubation, Staphylococcus aureus (Wood strain without protein A) BioParticles™, Alexa Fluor™ 488 conjugate (Invitrogen, ThermoFisher Scientific) were added to the cells at a ratio of 1:1 (cells to particles). The cells and particles were incubated for 1 hour. At the end of incubation, cells were scraped from the wells, and the suspension spun down at 400 RCF for 4 min, the cell pellet was washed and suspended in a staining buffer. Cells were then stained with a monoclonal antibody against CD45 (clone 30-F11) and Ly6G (clone 1A8) for 30 min. at 4°C. Further, cells were re-suspended in PBS containing PI at a concentration of 2 µg/mL. Percent uptake of these particles by neutrophils (CD45^+^Ly6G^+^) was quantified using flow cytometry.

### Histopathology and Lung injury Scoring

For Histology analysis, the left lung lobe was dissected and fixed using 10% buffered formalin and embedded into paraffin for tissue sectioning. 5 µm sections were stained with hematoxylin-eosin (H&E). All H&E-stained slides were coded and observed under a binocular optical microscope (Olympus CX43). To prevent bias, the slides were coded, and the pathologist was blinded to the groups. A weighted histology grading scheme consistent with criteria outlined by the American Thoracic Society for animal models with ARDS was used for scoring^60^. Briefly, twenty random non-overlapping fields (at least 50% of each field was occupied by lung alveoli) were captured from each animal lung section and evaluated for the following parameters: neutrophils in the alveolar space, neutrophils in the interstitium, hyaline membrane formation, proteinaceous debris filling airspace, and alveolar septal thickening. Every variable received a score of 0, 1, or 2 based on its severity, with varying weights assigned according to the importance of these variables. The total of the weighted variables was calculated based on the number of identified fields, resulting in a final score ranging from 0 to 1.

### Lung stiffness measurement using micro indentation technique

For lung stiffness measurement using atomic force microscopy (AFM), the right inferior lobe was dissected and mounted onto 3% agarose gel within a 50mm petri dish. The gel volume is optimized such that the bottom portion of the lobe solidifies with the agar to immobilize tissue onto the petri dish. The petri dish with the lung lobe is filled with 1x PBS and subjected to wet-mode AFM. The cantilever used for FD spectroscopy is HYDRA6V-200NG with a 5 µm diameter SiO₂ sphere attached to it. Force constant of the cantilever is 0.045 N/m. The AFM used for the measurements is the Park Systems XE-Bio AFM. Additionally, the indentation depth was 1 µm.

Twenty random indentations were performed per lung lobe (or per animal), and an FD curve generated per indentation was converted to point data and processed through a modified version of a previously used automated MATLAB code^61^ for quality control and to measure Young’s modulus. The code is available here - https://github.com/Immunoengineeringlab/AFM_FD-curve-analysis-using-MATLAB.git.

Depending on the number of animals per experimental condition, the histogram of young modulus distribution was made, and distribution fit was performed by probability distribution fit on MATLAB.

### Lung hydroxyproline content

The right inferior lung lobe was retrieved from agarose gel post-AFM analysis and was homogenized by the flash-freezing method and lyophilized for 48 hours. The lyophilized sample was subjected to acid hydrolysis and resuspended in deionized water. The samples were assayed for hydroxyproline using chloramine-T, as described^62^.

### rmG-CSF model

To mobilize immature neutrophils into peripheral blood, mice were injected (intraperitoneally, i.p.) with mouse G-CSF Recombinant protein (100ug/Kg, PeproTech, Israel) 72 hours and 48 hours prior to LPS administration.

### Treatment with small molecules and neutralizing antibodies

Mice were either treated with drugs and monoclonal antibodies of interest at different time points and doses depending on the study design. The dose of sivelestat sodium salt (50mg/Kg, Tocris Biosciences), and vehicle control (saline) were optimized and injected via i.p. for 5 consecutive days from the time of LPS administration. Isotype control (Bioxcell), anti-G-CSF (Invitrogen), anti-IL1β (Bioxcell) and anti-TNFα (Bioxcell) were optimized and injected i.p., 24 hrs. prior, and 24 hrs. and 72 hrs. post-LPS administration (3 doses). The effects of all the therapeutics were assessed 14-day post LPS administration.

### Single-cell isolation, library construction, and sequencing

Immune cells from blood and BALF were isolated as described in earlier sections. Sample tagging was performed by labeling cells with BD® Mouse Single-Cell Multiplexing Kit. Neutrophils in the samples were tagged with Ly6G antibody-derived tags (Ly6G-ADTs, BD® AbSeq Ab-Oligos). These cells were subjected to single cell capture and cDNA Synthesis, library construction using a Whole Transcriptome Analysis (WTA, BD) kit with the BD Rhapsody™ single-cell analysis system following standard procedures provided by the manufacturer. Sequencing was performed by Eurofin Advinus (Bengaluru, India). The sequenced data and the codes used for analysis are available here – https://github.com/Immunoengineeringlab/ScRNA-Seq_Neutrophils_2025_VKD_AS.git.

### Single-cell RNA seq data processing and cell type annotation

Pre-processing of the raw FASTQ files obtained from the BD Rhapsody™ single-cell analysis system, including read quality filtering, alignment, and quantification, was performed on the Seven Bridges platform using the BD Rhapsody™ sequence analysis pipeline with default references including RhapRef_Mouse_WTA_2023-02 for mRNA and Mouseneutrophil_Ly6G for AbSeq. Further, downstream analyses were performed using R 4.4.1 and Seurat 5.1.0^63^. Cells marked "Undetermined" or "Multiplet" by BD Rhapsody™ Sequence Analysis Pipeline, cells failing to have a minimum of 300 genes (min.features = 300), and genes not present in at least three cells (min.cells = 3) were filtered out. To retain only high-quality cells, cells with nFeature_RNA in the range 500-4500 and 500-8000, nCount_RNA in the range 1000-25000 and 1000-50000 for blood and BALF, respectively, and mitochondrial percentage (percent.mt) < 15 in all groups were used for downstream analyses. This filtering led to a total of 9843 cells from the single inflammation model of blood (3506 mock-treated with an average of 2560 genes/cell, 6337 chitosan implanted with an average of 2601 genes/cell), and 6843 (2577 mock-treated with an average of 2132 genes/cell, 2075 chitosan implanted with an average of 2075 genes/cell) and 9772 (5548 mock-treated with an average of 3302 genes/cell, 4224 chitosan implated with an average of 3695 genes/cell) cells from the double inflammation model of blood and BALF, respectively.

The cell types were annotated using SingleR^64^ against the ImmGen dataset accessed through the celldex package, while the cell proportions in each Seurat object were quantified and visualized using the scMega 1.0.2 package. Blood and BALF white blood cells (WBCs) and their cell types were analyzed independently.

### Batch correction, Dimensional reduction, and Unsupervised clustering

Data analysis was performed using standard Seurat integration with SCTransform-normalized dataset workflow. In brief, integration was performed using the CCA Integration method to account for batch effects and any confounding factors between mock and chitosan implanted SCTransform-normalized data (with mitochondrial mapping percentage regression). The number of dimensions for dimensional reduction to run FindNeighours() and RunUMAP() was chosen as the minimum value between the principal component (PC), which exhibits cumulative percent > 90%, and % variation associated with the PC as < 5, and the last PC where the change of % of the variation with the subsequent PC is more than 0.1%. A resolution of 0.5, 0.2, and 0.4 was used for clustering using FindClusters() for the WBCs (blood and BALF) and the Neutrophil subsets of blood and BALF, respectively, and clusters were visualized using Uniform Manifold Approximation and Projection (UMAP).

To analyze neutrophils with high protein expression, neutrophils from the WBC clusters were further subset to retain neutrophils with a median expression of Ly6G-ADT > 50% and analyzed independently. Further, to analyze immature neutrophils from this subset, neutrophils with no Cd101 expression were also separately analyzed.

### Identification of differentially expressed genes (DEGs) and their functional significance

The PrepSCTFindMarkers() function was run on the SCtransformed counts, after which genes with a log2 Fold change of 1.5 and detected in a minimum of 30% of cells in chitosan implanted compared to mock-treated cells with adjusted p-value < 0.05 as identified by the FindMarkers() function were considered DEGs. DEGs were identified for each cell type. Further, the most significantly perturbed markers for each Seurat cluster compared to the other clusters were also identified using the FindAllMarkers() function using logfc.threshold = log2(1.5) and min.pct = 0.60.

The functional significance of all markers was identified and visualized using the clusterProfiler 4.12.2 package against the Gene Ontology (GO) and KEGG databases. The GO biological processes were further filtered with the simplify() function with a cut-off of 0.7 to remove some redundant terms.

### Scoring of biological process

Scoring of various biological processes of high relevance to neutrophils, including NET formation, neutrophil maturation, neutrophil extravasation, azurophilic granule, neutrophil specific granule, neutrophil gelatinase granule, neutrophil secretory granule, neutrophil aging, positive regulation of apoptosis, phagocytosis, chemotaxis, neutrophil activation, NADPH Oxidase, and ROS were done against previously curated gene signatures from Xie et al.^29^ using the AddModuleScore() function. Additionally, we curated gene signatures for apoptotic neutrophils uptake induced gene score by macrophages from Liebold et al.^33^ Resident airspace macrophage (ResAM) and Recruited airspace macrophage (RecAM) gene signatures from Mould et al.^65^ Efferocytosis receptors gene module score was curated by us (Axl, Mertk and Tyro3). The statistical differential scoring of the processes between chitosan and mock-treated cells was quantified using the Wilcoxon rank sum test.

### Correlation of scRNA-seq-defined neutrophil populations with previously reported neutrophil sub-populations

The similarity between clusters identified in our study and previously reported neutrophil subpopulations^29^ was quantified using pairwise Jaccard similarity among their respective top 30 cluster marker labels. The associated p-value was determined by permutation hypothesis testing with 1000 iterations of cluster marker labels.

Further, FDR-corrected p-values were also assigned to each observed similarity value. Clusters with adjusted p-values < 0.05 were considered similar.

### Cell communication

CellChat 1.6.1 was used to identify major signaling changes across cell types and their communication in chitosan and mock-treated cells. The BALF WBC Seurat object was first split into mock and chitosan implanted, then individually converted to CellChat objects and analyzed using the CellChatDB.mouse database. The expression data of a subset of genes involved in signaling was taken, after which the overexpressed genes and interactions in both the Cellchat objects were identified, and communication probabilities were computed. Cell-cell communication was further filtered at a threshold of min.cells = 10, and cell-cell communication at a signaling pathway level was inferred. Aggregated cell-cell communication networks and network centrality scores using the “netP” slot were also quantified. Signaling from each cell type was visualized using circle plots, and cell-cell communication in pathways such as CSF, IL1 and TNFα signaling was visualized using chord plots.

### Statistical Analysis

For data that was normally distributed or expected to be normally distributed, Student’s ‘t’ test (for comparison between 2 groups), one-way ANOVA (for comparison across multiple groups with one variable) or two-way ANOVA (for comparison across multiple groups with two variables) followed by a post-hoc test was used. When data was analyzed using one of the above statistical tests, significance is denoted using the * symbol as explained in the figure legends. When the data was not normally distributed or distribution was unknown, the Wilcoxon rank sum test has been used. Exact p values from the Wilcoxon tests are reported in the figures.

## Acknowledgements

This research was funded by the DBT/Wellcome Trust India Alliance Intermediate fellowship to SJ (IA/I/19/1/504265). AA was supported by a fellowship from the Department of Biotechnology, Govt. of India. VKD and RG are supported by the Prime Minister’s Research Fellowship, Govt. of India. The authors acknowledge Sreenath Balakrishnan for providing the initial codes to analyze the atomic force microscopy data. The authors thank the central animal facility at the Indian Institute of Science (IISc) for their support. Bridging financial support from the IISc is also acknowledged.

