## Supplementary for "Chronic Inflammation Induced Immature Neutrophils Drive Immunopathology During Subsequent Inflammatory Events"

**Table of Contents:**

Supplementary Figure 1 – 8 (pages 2 – 15)

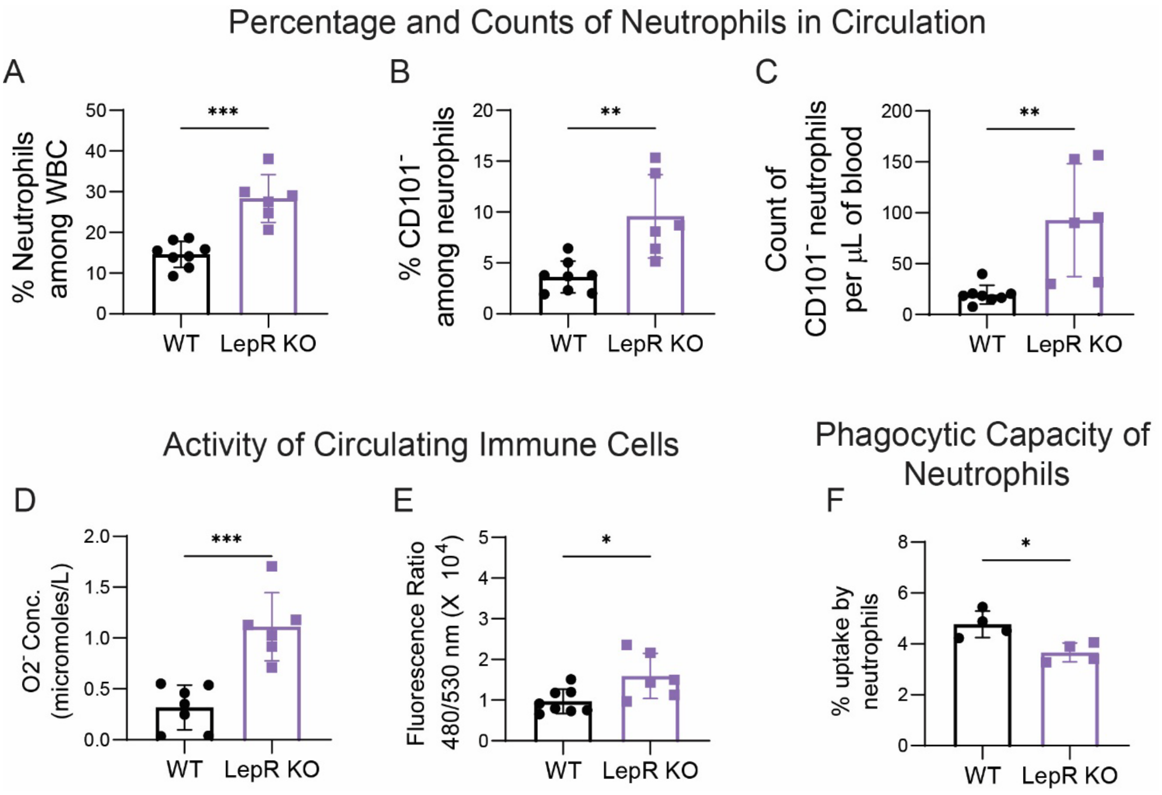


**Supplementary Figure 1: Counts and function of circulating neutrophils in obese-diabetic leptin receptor knockout (LepR KO) mice.**

**A-C –** Flow cytometry-based quantification of percentage of total neutrophils among white blood cells (WBCs) in circulation (A), percentage of immature neutrophils among total neutrophils (B), and total number of immature neutrophils per microliter of blood (C). n=8 for WT and 6 for LepR KO.

**D and E –** Extracellular reactive oxygen species (ROS) quantified using cytochrome C assay (D) and extracellular DNA quantified using sytox assay (E) measured by ex vivo analysis of WBCs obtained from these mice. n=7 or 8 for WT and n=6 for LepR KO.

**F –** Phagocytosis measured as percent uptake of A488-conjugated S. aureus bioparticles by circulating neutrophils in an ex vivo assay quantified using flow cytometry. n=4 for WT and n=4 for LepR KO.

All samples were biologically independent, and values presented are from three or more independent experiments. For statistical analysis, Student’s t test was used with * representing $p<0.05$, ** $p<0.01$, and *** $p<0.001$.

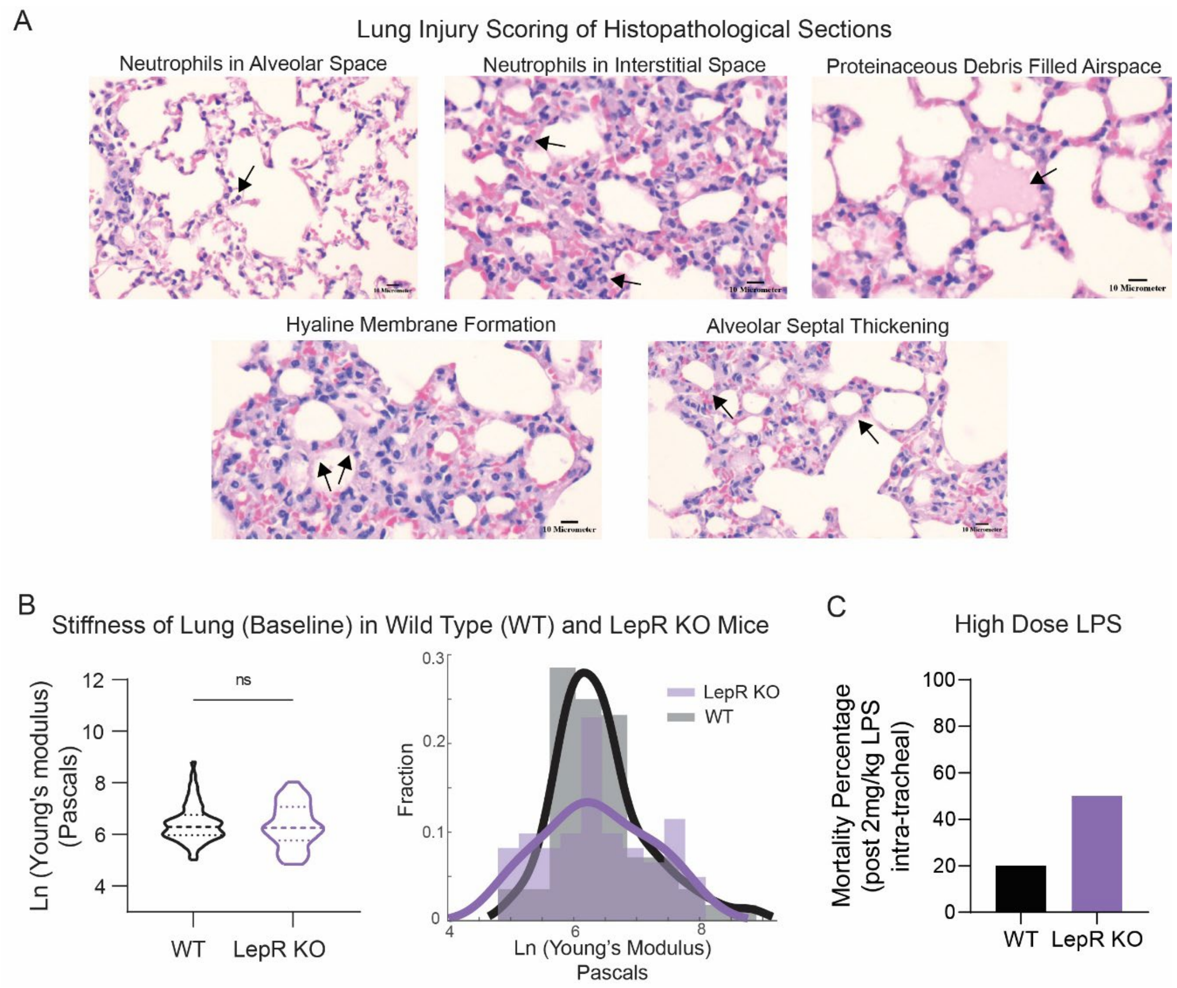


**Supplementary Figure 2: Lung injury scoring, lung stiffness at baseline levels and effect of high dose of lipopolysaccharide (LPS).**

**A –** Representative images of parameters used in scoring system for quantification of “overall” lung injury score. scale bar = 10μm

**B –** Lung stiffness measured by micro-indentation using atomic force microscopy (AFM) (left) and distribution profile of Young’s modulus obtained from each measurement (dark lines indicate curves fitted on the histograms using a probability distribution fit). n=59 for WT and n=61 for LepR KO micro indentation points sampled from n≥3 animals.

**C –** Percentage of mortality observed post-LPS administration with the dose of 2mg/kg. n=20 animals per group.

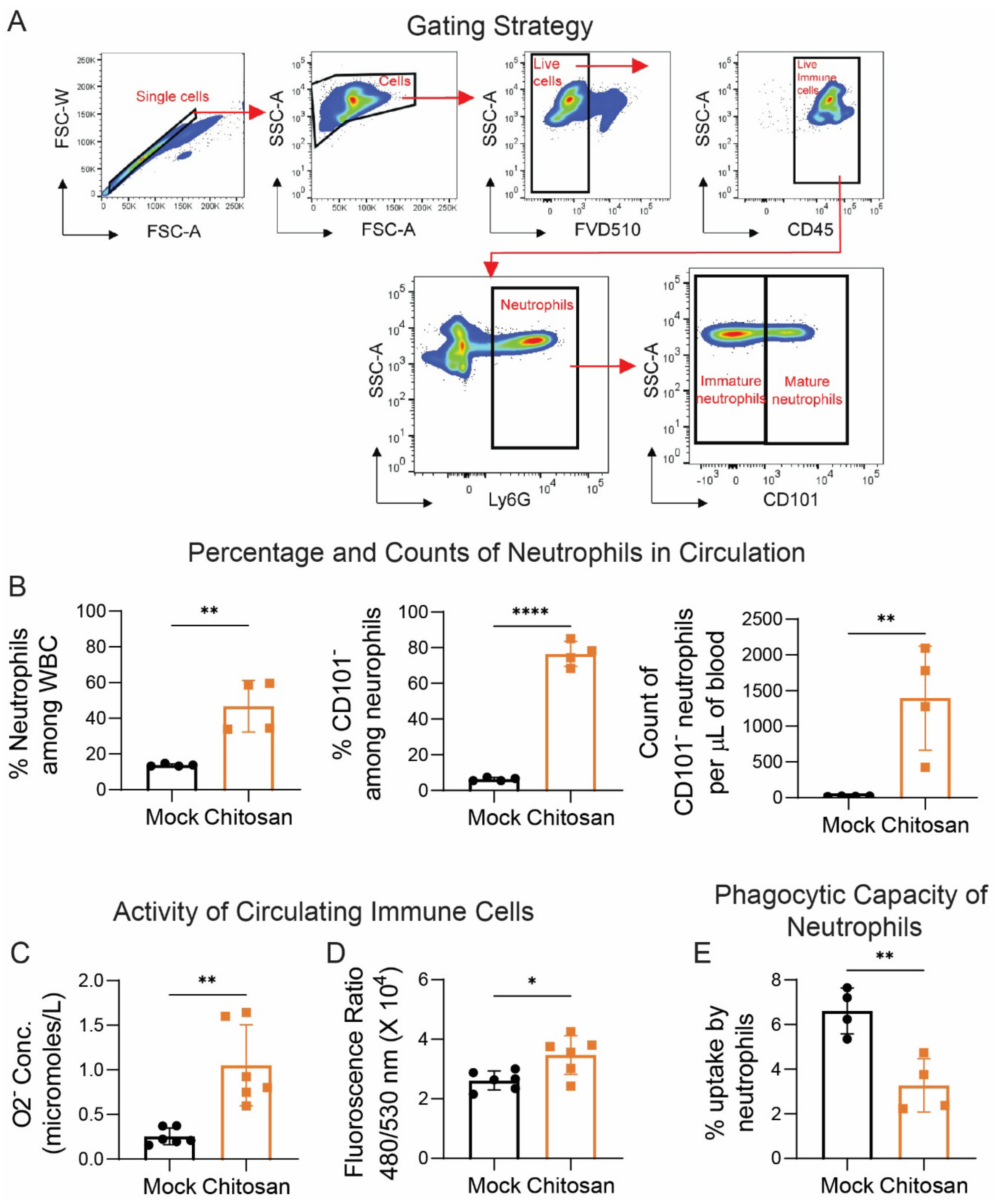


**Supplementary Figure 3: Counts and function of circulating neutrophils in mock-surgery and chitosan-implanted mice.**

**A –** Representative flow cytometry dot plots and gating strategy to identify immature and mature neutrophils.

**B –** Flow cytometry-based quantification of percentage of total neutrophils among white blood cells (WBCs) in circulation, percentage of immature neutrophils among total neutrophils, and total number of immature neutrophils per microliter of blood. n=4 animals per group.

**C and D –** Extracellular reactive oxygen species (ROS) quantified using cytochrome C assay (C) and extracellular DNA quantified using sytox assay (D) measured by ex vivo analysis WBCs obtained from these mice. n=6 animals/group.

**E –** Phagocytosis measured as percent uptake of A488-conjugated S. aureus bioparticles by circulating neutrophils in an ex vivo assay quantified using flow cytometry. n=4 animals per group.

All samples were biologically independent, and values presented are from three or more independent experiments. For statistical analysis, Student’s t test was used with * representing $p<0.05$, ** $p<0.01$, and **** $p<0.0001$.

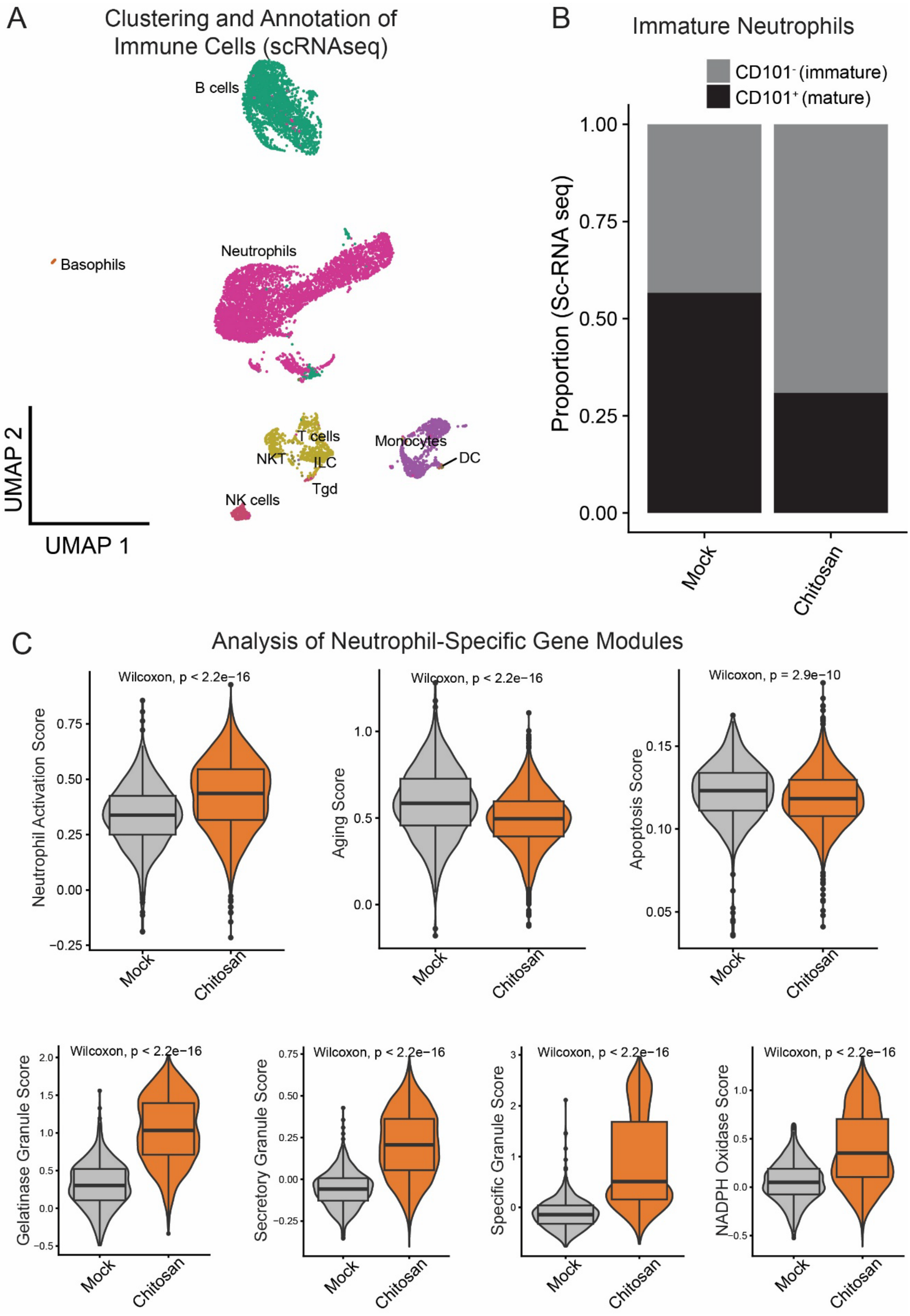


**Supplementary Figure 4: Analysis of single cell RNA sequencing (scRNAseq) data.**

**A –** Unbiased uniform manifold approximation and projection (UMAP) of WBCs in circulation (10-days post chitosan implantation). Each dot represents an individual cell and is colored according to cell cluster.

**B –** Proportion of mature/immature neutrophils from the scRNAseq data.

**C –** Comparison of neutrophil-specific functional scores based on the expression of groups of genes contributing to this function. The significance among functional scores was determined using the Wilcoxon test.

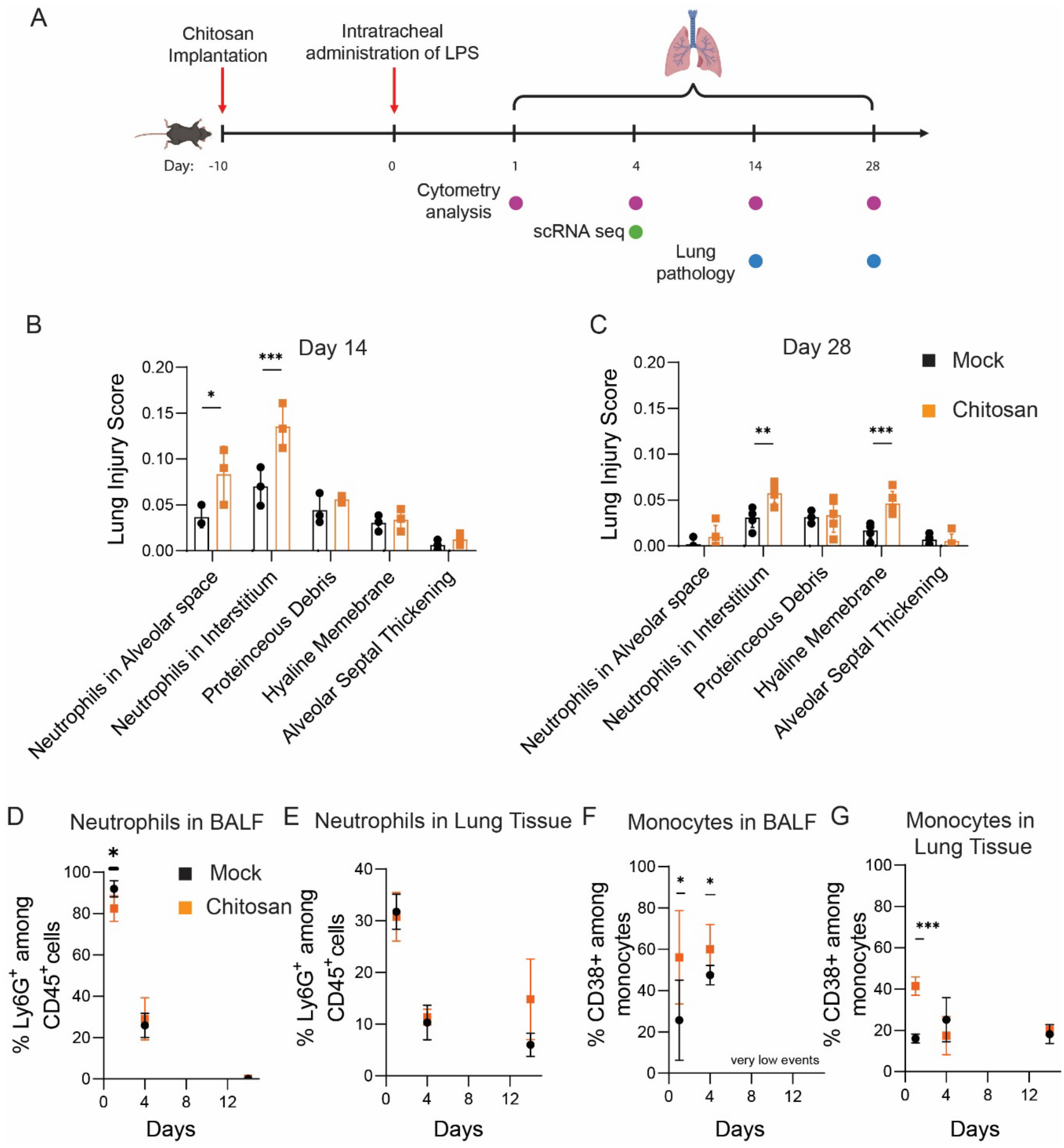


**Supplementary Figure 5: LPS-mediated sterile lung injury model in chitosan-implanted mice.**

**A –** Schematic of study design.

**B and C** – Lung damage was quantified from histology images and represented as a sum of individual weighted scores of each parameter at day 14 (B, n=3 animals/group) and day 28 (C) post-LPS administration (n=5 animals/group).

**D – G** – Quantification of the kinetics of cell proportions in the bronchoalveolar lavage fluid (BALF) or ling tissue, with **D** representing the percentage of neutrophils among in BALF, **E** representing neutrophils in in lung tissue, **F** representing percentage of $CD38^+$ monocytes among total monocytes in BALF, and **G** representing these monocytes in lung tissue (n≥3 animals/group/timepoint).

All samples were biologically independent, and values presented are from three or more independent experiments. For statistical analysis, Student's t test, or one-way or two-way ANOVA followed by appropriate post-hoc test was used with * representing $p<0.05$, ** $p<0.01$, *** $p<0.001$, and **** $p<0.0001$.

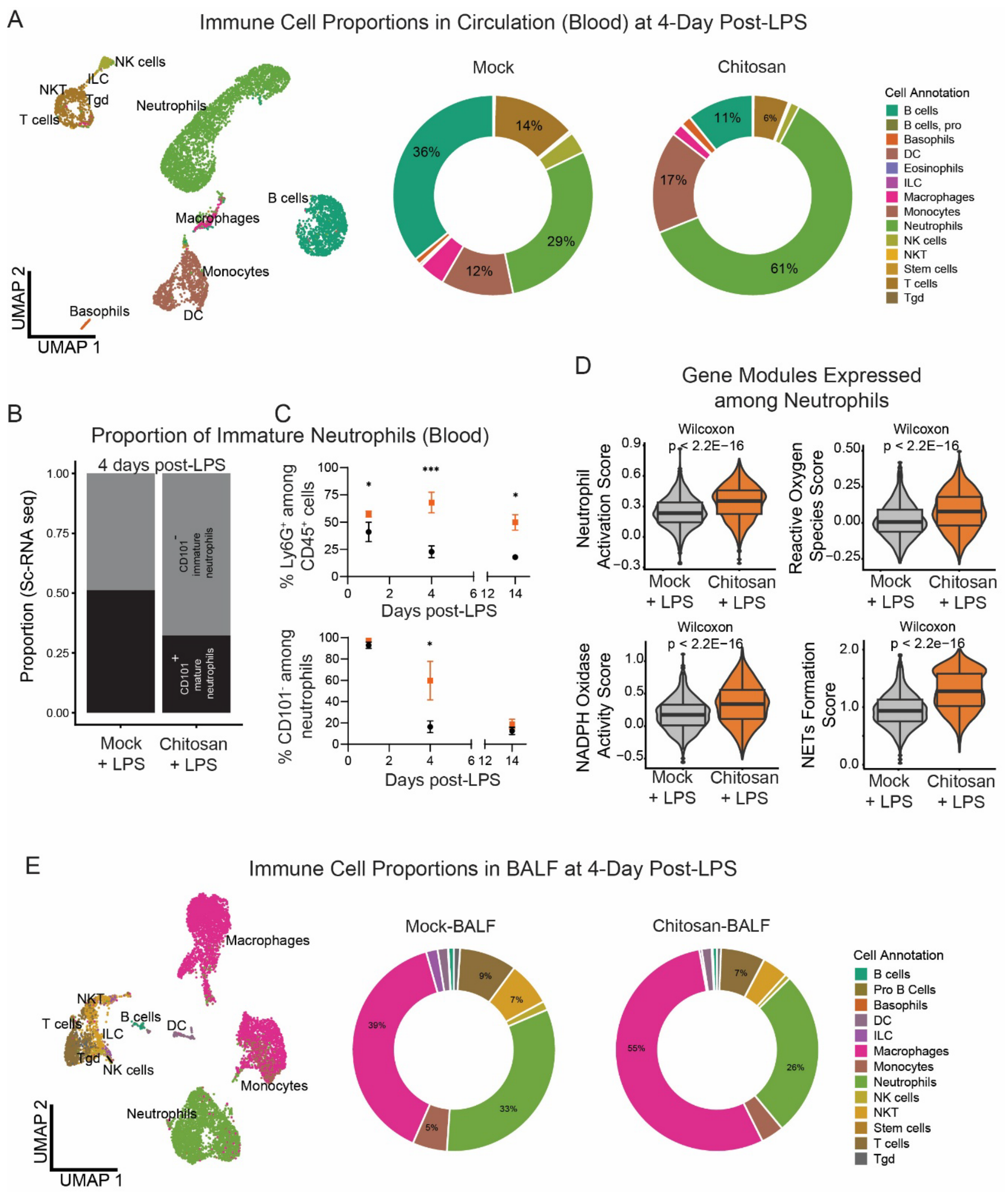


**Supplementary Figure 6: single cell RNA sequencing (scRNAseq) analysis of cells obtained from circulation and bronchoalveolar lavage fluid (BALF) at 4-days post LPS administration in chitosan implanted mice and respective controls.**

**A –** Unbiased uniform manifold approximation and projection (UMAP) of WBCs in circulation (4 days post LPS administration) and their proportions represented in a donut plot. Each dot represents an individual cell and is colored according to cell cluster.

**B –** Proportion of mature/immature neutrophils from the scRNAseq data.

**C –** Percentage of neutrophils and immature neutrophils quantified at 1-, 4- and 14-days post LPS administration using flow cytometry (n≥3 animals/group).

**D –** Neutrophil-specific functional scores of circulating neutrophils (4 days post LPS administration) based on the expression of groups of genes contributing to this function. The significance among functional scores was determined using the Wilcoxon test.

**E –** Unbiased uniform manifold approximation and projection (UMAP) of WBCs in BALF (4 days post LPS administration) and their proportions in donut plot. Each dot represents an individual cell and is colored according to cell cluster.

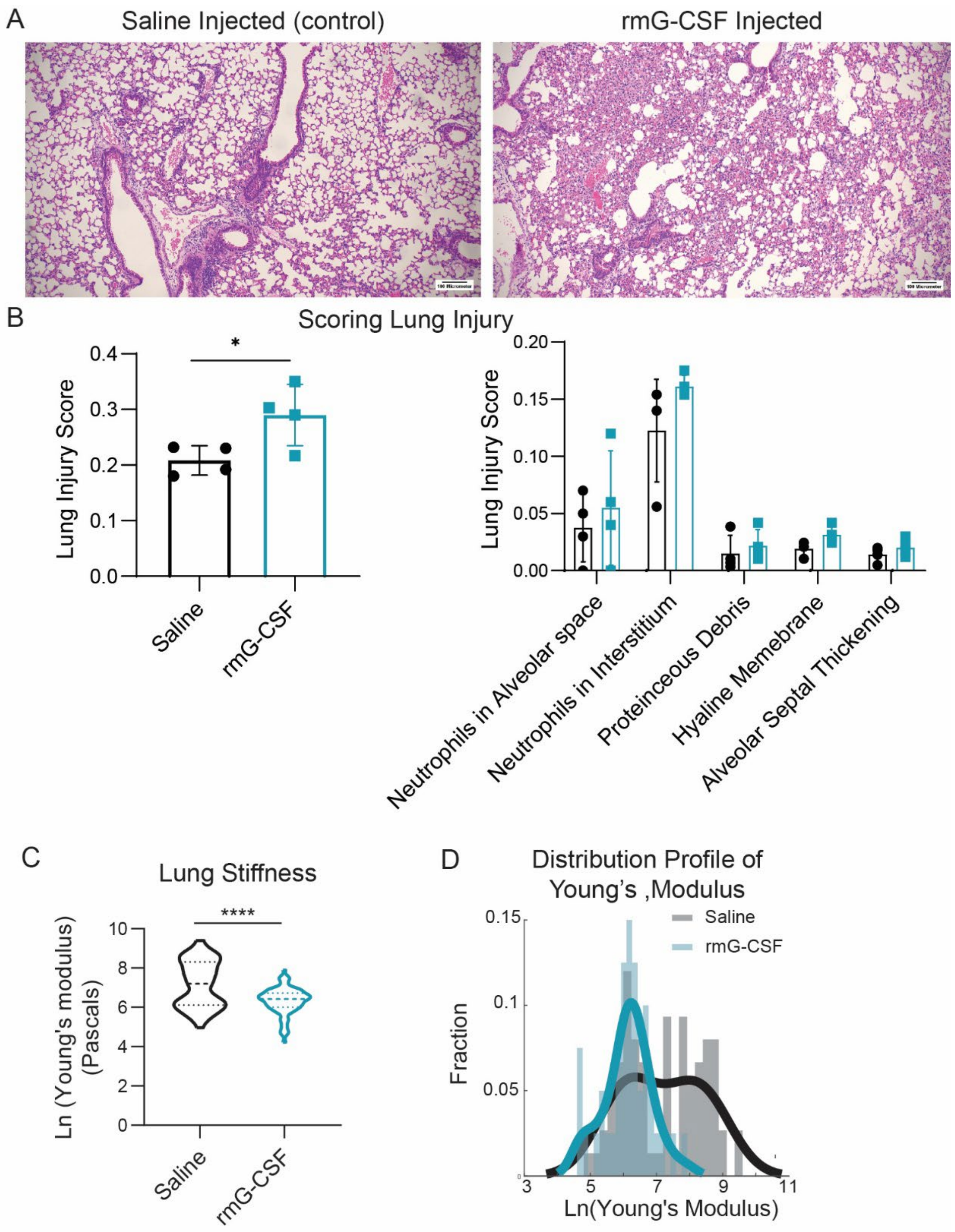


**Supplementary Figure 7: Recombinant murine G-CSF (rmG-CSF) administration.**

**A –** Representative images of hematoxylin and eosin-stained tissue sections at 14 days post LPS administration in saline vs recombinant G-CSF injected mice. scale bar = 100μm.

**B –** Lung damage was quantified from these images and represented as overall injury score and a breakdown of individual weighted scores from each parameter used for overall scoring of the images. n=4 animals/group.

**C and D –** Lung stiffness measured by micro-indentation using atomic force microscopy (C), and the distribution profile of Young's modulus (D)

obtained from each measurement (dark lines indicate curves fitted on the histograms using a probability distribution fit). For saline-injected (N=77) and recombinant murine G-CSF injected mice (N=82); (N-represents number of micro indentations sampled across n=4 animals/group).

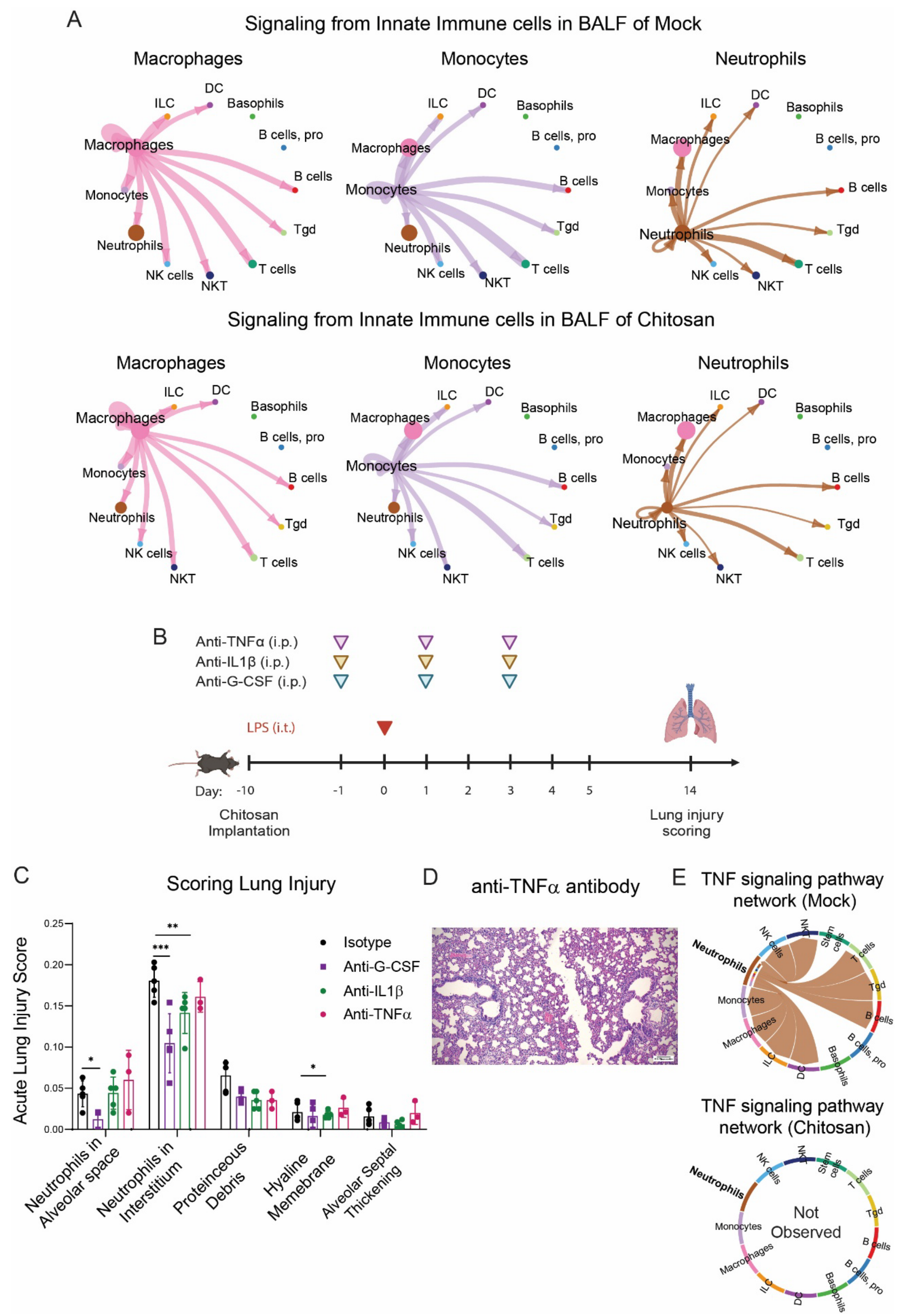


**Supplementary Figure 8: Cellchat-based analysis of signaling networks and associated studies involving blocking of specific proteins using antibodies.**

**A –** Cross-talk among immune cells quantified by CellChat using our scRNAseq data from bronchoalveolar lavage fluid (BALF) of mock and chitosan-implanted mice, with a focus of communication initiated by macrophages (left), monocytes (middle) and neutrophils (right).

**B –** Schematic of study design.

**C –** Lung damage was quantified from histology images and represented as a breakdown of individual weighted scores from each parameter for chitosa- implanted mice treated with isotype control (n=5), anti-GCSF(n=5), anti-IL1β (n=5) and anti-TNFα (n=3); 14 days post LPS administration.

**D –** Representative images of hematoxylin and eosin-stained tissue sections following treatment with anti-TNFα. Scale bar = 100μm.

**E** – Chord plot of TNFα signaling network analysis using Cellchat performed on immune cells of BALF (4 days post LPS administration).
